# The LC3B FRET biosensor monitors the modes of action of ATG4B during autophagy in living cells

**DOI:** 10.1101/2022.05.06.490917

**Authors:** Elif Begüm Gökerküçük, Angélique Chéron, Marc Tramier, Giulia Bertolin

## Abstract

Although several mechanisms of autophagy have been dissected in the last decade, following this pathway in real time remains challenging. Among the early events leading to its activation, the ATG4B protease primes the key autophagy player LC3B. Given the lack of reporters to follow this event in living cells, we developed a Förster’s Resonance Energy Transfer (FRET) biosensor responding to the priming of LC3B by ATG4B. The biosensor was generated by flanking LC3B within a pH-resistant donor-acceptor FRET pair, Aquamarine/tdLanYFP. We here showed that the biosensor has a dual readout. First, FRET indicates the priming of LC3B by ATG4B and the resolution of the FRET image allows to characterize the spatial heterogeneity of the priming activity. Second, quantifying the number of Aquamarine-LC3B puncta determines the degree of autophagy activation. We then showed that there are pools of unprimed LC3B upon *ATG4B* downregulation, and that the priming of the biosensor is abolished in *ATG4B* knockout cells. The lack of priming can be rescued with the wild-type ATG4B or with the partially active W142A mutant, but not with the catalytically dead C74S mutant. Moreover, we screened for commercially-available ATG4B inhibitors, and we illustrated their differential mode of action by implementing a spatially-resolved, broad-to-sensitive analysis pipeline combining FRET and the quantification of autophagic puncta. Finally, we uncovered the CDK1-dependent regulation of the ATG4B-LC3B axis at mitosis. Therefore, the LC3B FRET biosensor paves the way for a highly-quantitative monitoring of the ATG4B activity in living cells and in real time, with unprecedented spatiotemporal resolution.

## Introduction

Conserved in all eukaryotic cells, macroautophagy/autophagy is the lysosome-mediated degradation and recycling of the intracellular components [1]. Autophagy is triggered as a survival response in paradigms such as starvation, pathogen infection or DNA damage, and it contributes to cellular differentiation, immunity, aging and cell death [2–4]. In mammals, autophagy starts at sites of endoplasmic reticulum (ER) enriched for phosphatidylinositol 3-phosphate [PI(3)P]. On these subdomains, a double-membrane structure termed phagophore forms [5]. As the phagophore elongates into a crescent-shaped structure, it engulfs bulk or selective cargoes and then closes into a double-membrane vesicle, the autophagosome. The fusion of autophagosomes with lysosomes results in the degradation of the sequestered cargo by the lysosomal acid hydrolases.

A series of AuTophaGy-related (ATG) proteins regulate the autophagic pathway [6]. Among them, a special attention is given to the ATG8 family, which are the key proteins found on autophagosomal membranes at all stages of the pathway. ATG8 proteins are ubiquitin-like adaptor proteins involved in autophagosome formation, biogenesis and cargo selection [7–9]. In mammals, ATG8 proteins belong to two subfamilies, the MAP1LC3/LC3 (microtubule-associated protein 1 light chain 3) and GABARAP (gamma-aminobutyric acid [GABA] type A receptor-associated protein) [10,11]. A total of seven genes - *LC3A, LC3B, LC3B2, LC3C, GABARAP, GABARAPL1* and *GABARAPL2* - code for the LC3 and GABARAP subfamilies in humans [10]. LC3/GABARAPs are found as inactive pro- forms upon translation, and are activated by the ATG4 family of cysteine proteases [12,13]. In humans, four members of the ATG4 family (ATG4A, B, C and D) are responsible for this activation step, which is to proteolytically cleave the C-terminus of pro-LC3/GABARAP proteins and convert them into the so-called form-I. This crucial cleavage is known as “pro-LC3/GABARAP priming”, and it is essential to expose a specific glycine residue required for the lipidation of the cytosolic LC3/GABARAP-I proteins to the phosphatidylethanolamine (PE) head groups of the forming phagophores. This is achieved after a series of reactions that involves the E1-like enzyme ATG7, the E2-like enzyme ATG3 and the E3-like complex ATG12–ATG5-ATG16L1 [12,14–16]. The PE-conjugated LC3/GABARAP proteins are then called LC3/GABARAP-II, and function in membrane tethering, hemifusion, autophagosome formation and cargo recruitment [17–20]. Once the phagophore is fully closed, LC3/GABARAP-II proteins are removed from the outer surface of the phagophore membrane by ATG4s, through the hydrolysis of the link between PE and LC3/GABARAP [12]. Although the importance of this second round of cleavage activity (referred as deconjugation hereafter) by ATG4 was shown to be important for the normal progression of autophagy in yeast [21–23], recent studies in human cells suggest the existence of autophagy-independent roles for the deconjugation activity of ATG4s [24,25]. Therefore, the relevance of ATG4-mediated deconjugation for the progression and completion of autophagy in models other than yeast still requires further investigation [21–26].

Autophagy plays an essential role to maintain cellular homeostasis, and its dysfunction has been implicated in many pathological conditions such as neurodegenerative diseases, cancer, inflammation, muscular and hearth disorders [27]. As a consequence, therapeutic options to modulate autophagy emerged as promising strategies for the treatment of these complex pathologies [28]. In this light, targeting ATG4s to inhibit autophagy in its early stages has a significant potential to completely block autophagy [29]. However, currently available compounds targeting ATG4 activity show poor specificity and/or efficacy [30]. In addition, there is a lack of dedicated probes that can be used in living cells to monitor ATG4 activity during autophagy progression. Overall, this creates a bottleneck for the identification of ATG4 inhibitors with improved properties. For these reasons, we developed a Förster’s Resonance Energy Transfer (FRET) biosensor, named the LC3B biosensor, to simultaneously monitor: 1) the priming of LC3B by ATG4 and 2) the accumulation of LC3B on the autophagic membranes.

The FRET phenomenon is a non-radiative energy transfer between a donor and an acceptor pair of fluorophores. FRET can occur when the emission spectrum of the donor fluorophore partially overlaps with the excitation spectrum of the acceptor, and this only when the two fluorescent moieties are in close proximity (less than 10 nm apart) [31]. This phenomenon can be used to monitor many different cellular events including the exploration of protein-protein interactions, the changes in conformation of proteins, and the up- or downregulation of signaling pathways [32,33]. With the recent advances, FRET quantification by fluorescence lifetime imaging microscopy (FLIM) became a very useful method to study molecular activities in living cells [34].

In this study, we present the LC3B biosensor as a superior probe that can be used in living cells to monitor the activation – LC3B priming by ATG4 – and progression – LC3B accumulation on the autophagic membranes – of autophagy, in real time and with high spatial resolution. We show that the biosensor recapitulates the main features of the endogenous LC3B protein in terms of forming puncta-shaped structures, of ATG4B-dependent cleavage, and of its colocalization with lysosomal proteins upon autophagy induction and/or lysosomal inhibition. We also show that the biosensor can report on the changes in proLC3B priming, and this in an ATG4B-dependent manner. Using *ATG4* knockout cells, we demonstrate that the absence of ATG4B maximizes the FRET response of the biosensor as a consequence of the complete lack of proLC3B priming. We then show that proLC3B priming can be rescued with the ectopic expression of the wild-type ATG4B. Interestingly, we demonstrate that the ATG4B^W142A^ mutant, previously shown to possess a significantly reduced catalytic activity [35], is capable of rescuing proLC3B priming similarly to the wild-type protein. By using the LC3B biosensor and performing multiple approaches to analyze FRET/FLIM, we report on the action of mechanisms of available ATG4 inhibitors. By doing so, we provide a framework of how to use the LC3B biosensor and analyze the acquired data to identify new ATG4 inhibitors with better specificity and efficacy. Finally, we used the biosensor to reveal the involvement of the cell cycle protein CDK1 in the ATG4B-LC3B axis at mitosis, a cell cycle phase where the involvement of autophagy is still controversial.

## Results

### The LC3B biosensor dynamically reports on the activation or the inhibition of the autophagy flux, and it colocalizes with LAMP2 in an autophagy-dependent manner

To monitor the priming activity of ATG4 in real time and with spatial resolution, we developed a FRET-based biosensor that can be utilized in living cells. We chose LC3B as it is a known target of the ATG4 activity that undergoes an ATG4-mediated cleavage on Gly120, and it is among the best characterized players of the autophagy pathway [36]. The biosensor was designed to flank the N- and C-termini of proLC3B with a donor-acceptor FRET pair (Fig. 1A). The FRET pair composed of Aquamarine (donor, cyan) and tdLanYFP (acceptor, yellow) was selected on the resistance of both fluorophores to acidic pH [37–39]. In the absence of ATG4 activity, the LC3B biosensor is expected to remain unprocessed in cells, allowing Aquamarine and tdLanYFP to perform FRET (Fig. 1A). If ATG4 is active, the biosensor is expected to undergo an ATG4-dependent proteolytic cleavage at its C-terminus, thereby losing the tdLanYFP moiety and the FRET effect with it. This allows to follow the initial C-terminal priming activity of ATG4 as an early, FRET-based readout. In addition, the priming of LC3B leads to its conversion into the I form, which will still be tagged by Aquamarine (Aquamarine-LC3B-I). When the resulting Aquamarine-LC3B-I protein is integrated into the PE head groups of the phagophores, the biosensor is expected to function as a canonical fluorescent probe to quantify LC3B-positive puncta structures. Therefore, our biosensor also provides a quantitative readout on the late stages of the autophagic pathway, and it can be used to estimate the number of autophagosomes in individual cells (Fig. 1A).

**Figure 1.**
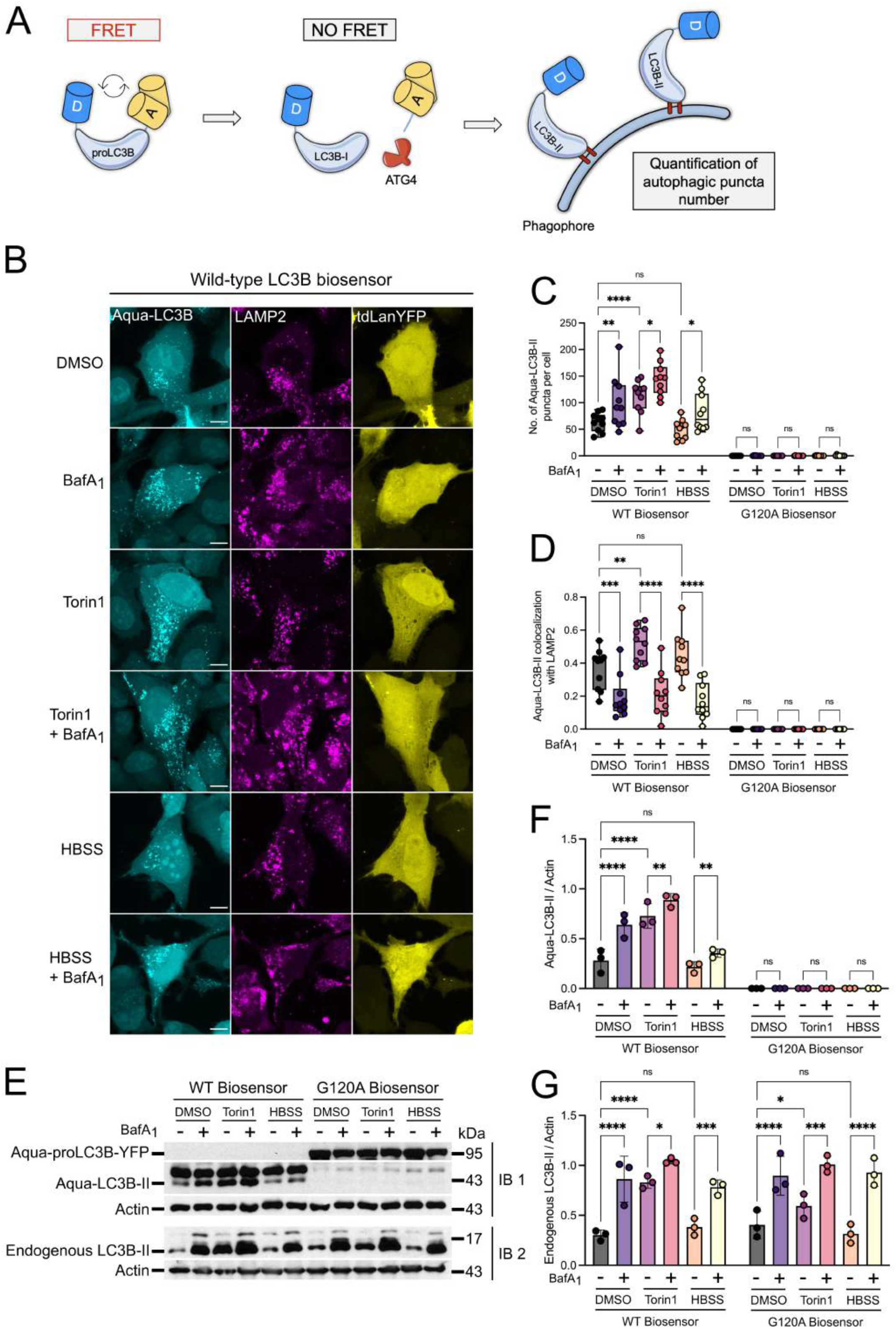
The LC3B biosensor reports on autophagy induction and/or lysosomal inhibition, and colocalizes with LAMP2 in an autophagy-dependent manner. (**A**) The cartoon illustrates the design and the mode of action of the LC3B biosensor. The biosensor was designed to flank the N- and C-termini of proLC3B with a donor (D, Aquamarine)-acceptor (A, tdLanYFP) FRET pair. When ATG4 is not active, the biosensor is expected to remain unprocessed in cells, allowing Aquamarine and tdLanYFP to perform FRET. Upon the proteolytic activity of ATG4, the biosensor is expected to be cleaved at its C-terminus, in turn losing its tdLanYFP moiety and the FRET effect with it. A successful priming of the biosensor is expected to yield Aquamarine-LC3B-I, which can then be integrated into the PE head groups of the phagophores and observed as puncta-shaped structures. The resulting Aquamarine-LC3B-II puncta-shaped structures can then be quantified to estimate the number of autophagosomes. (**B**) Representative fluorescence images of U2OS cells expressing the WT biosensor and stained for endogenous LAMP2. To investigate the changes in Aqua-LC3B puncta numbers and their colocalization with LAMP2, cells were treated with the following compounds: DMSO (6h), BafA1 (6h, 100 nM), Torin1 (3h, 250 nM), Torin1 (3h, 250 nM) + BafA1 (6h, 100 nM), HBSS (1h), HBSS (1h) + BafA1 (6h, 100 nM). Scale bar: 9 μm. (**C**) Quantification of the number of Aqua-LC3B-II puncta in cells expressing the WT or G120A biosensor and treated as indicated. (**D**) Quantification of the ratio of Aqua-LC3B-II puncta structures colocalizing with LAMP2-positive objects in cells expressing the WT or G120A biosensor and treated as indicated. *n* = 10 cells per condition from one representative experiment (of three) in (**C**) and (**D**). (**E**) Representative western blotting images and corresponding quantifications (**F, G**) of total lysates from U2OS cells expressing the WT or G120A biosensor and treated as indicated. IB1 and IB2 correspond to the same lysates blotted for overexpressed (IB1) or endogenous (IB2) LC3B forms. Loading control: Actin. *n* = 3 independent experiments \**P* < 0.05, \*\**P* < 0.01, \*\*\**P* < 0.001, \*\*\*\**P* < 0.0001, ns (not significant) as determined by two-way ANOVA with Tukey’s multiple comparison test in (**C**) and (**D**), and with two-stage step-up method of Benjamini, Krieger and Yekutieli’s multiple comparison test to control the false discovery rate in (**F**) and (**G**).

We first explored whether the LC3B biosensor is capable of localizing to autophagosomes, which normally appear as puncta-shaped structures. In U2OS cells transiently transfected to express the biosensor, Aquamarine was observed to be present both in puncta-shaped structures, which are compatible with LC3B-II, and in the cytosol, which is compatible with Aqua-LC3B-I (Fig. 1B). To confirm that the puncta-shaped structures were autophagosomes, cells were treated with autophagy inducers as Torin1 or HBSS, in the presence or absence of the lysosomal inhibitor Bafilomycin A_1_ (BafA_1_). Compared to control cells, a significant increase in the number of Aquamarine-positive puncta was observed in cells expressing the wild-type LC3B biosensor (WT biosensor) and treated with BafA_1_ or Torin1 alone (Fig. 1B, C). A further increase in the number of Aquamarine-positive puncta was observed when cells were treated simultaneously with BafA_1_ and Torin1. This indicates that the puncta-shaped structures observed under these conditions are Aquamarine-LC3B-II (Aqua-LC3B-II)-positive autophagosomes, since they respond to autophagy induction and to lysosomal inhibition alone or in combination. Conversely, autophagy induction by starvation did not cause any increase in the number of puncta-shaped structures when compared to the control (Fig. 1B, C). However, when starvation with HBSS was coupled with BafA_1_ treatment, we observed a significant accumulation of puncta-shaped structures (Fig. 1B, C). In all conditions tested, tdLanYFP appeared to be diffused in the cytosol, thereby showing a dramatic difference compared to the distribution of Aquamarine into puncta-shaped structures. This strongly suggests that tdLanYFP is cleaved along with the C-terminal part of proLC3B, and therefore it cannot colocalize with Aquamarine on the puncta-shaped structures. To corroborate this observation, we explored the distribution of an uncleavable variant of the LC3B biosensor, which is mutated on Gly120 (hereby, G120A biosensor) and cannot be primed by ATG4 [36]. In cells expressing the G120A biosensor, both Aquamarine and tdLanYFP were exclusively diffused in the cytosol (Fig. S1), and they exhibited no puncta-shaped structures under any treatment (Fig. 1C and S1). The difference in puncta numbers between the WT and the G120A biosensors supports the notion that the WT construct is cleaved at the C-ter, and that it efficiently forms autophagosome-related puncta structures. In this light, we verified the cleavage profiles of the WT and G120A biosensors by western blotting. While the G120A biosensor had a molecular weight of ~95 kDa – corresponding to Aquamarine + proLC3B^G120A^ + tdLanYFP –, the WT biosensor was cleaved in all conditions tested and appeared as two bands at ~45kDa and ~43kDa. These bands were compatible with the molecular weight of Aquamarine + LC3B-I at ~45 kDa, and of Aquamarine + LC3B-II at ~43 kDa (Fig. 1E). Consistent with the quantifications of puncta numbers in cells expressing the WT biosensor (Fig. 1B, C), the levels of the Aqua-LC3B-II band increased upon BafA_1_ or Torin1 treatment, but remained unaltered upon starvation with HBSS (Fig. 1E, F). A further increase in Aqua-LC3B-II abundance was observed upon the co-treatment with BafA_1_ and Torin1 when compared to BafA_1_ or Torin1 alone (Fig. 1E, F). Similarly, the combination of HBSS and BafA_1_ increased the levels of Aqua-LC3B-II when compared to HBSS alone (Fig. 1E, F). These findings suggest a rather rapid degradation of Aqua-LC3B-II via lysosomal turnover in U2OS cells upon starvation with HBSS. This also suggests that the degradation of Aqua-LC3B-II can be slowed down when starvation is coupled with a late-stage lysosomal inhibitor such as BafA_1_. This was previously reported to occur in several other cell lines [24,40,41], and it substantiates the importance of measuring the autophagy flux in the absence or presence of lysosomal inhibitors. The differences observed in the levels of Aqua-LC3B-II were also observable at the level of the endogenous LC3B-II, confirming that the lipidation levels of the LC3B biosensor are similar to those of endogenous protein upon autophagy induction and/or lysosomal inhibition (Fig. 1E, G).

To explore whether a portion of Aqua-LC3B-II puncta structures are capable of colocalizing with lysosomes, we analyzed their juxtaposition with the lysosomal protein Lysosomal Associated Membrane Protein 2 (LAMP2). Compared to control cells, we found that the colocalization of Aqua-LC3B-II puncta structures with LAMP2-positive objects significantly increased when autophagy is induced with Torin1, but not with HBSS (Fig. 1B, D). As expected in cells expressing the WT biosensor, we observed that treatment with BafA_1_ significantly reduced the colocalization of Aqua-LC3B-II puncta with LAMP2 compared to untreated, Torin1- or HBSS-treated cells. Conversely, the G120A biosensor did not colocalize with LAMP2 in any condition (Fig. 1D and S1). Overall, these results show that the WT biosensor colocalizes with LAMP2 in an autophagy-dependent manner.

Taken together, these results demonstrate that the LC3B biosensor is efficiently cleaved. The biosensor is capable of forming puncta structures that are consistent with the lipidated, Aqua-LC3B-II form, and they colocalize with the lysosomal protein LAMP2. The autophagy-dependent changes in the number of puncta-shaped structures or their degree of colocalization with LAMP2 indicate that the biosensor is capable of successfully reporting on autophagy induction and/or lysosomal inhibition.

### The LC3B biosensor responds to the changes in proLC3B priming in an ATG4B-dependent manner

After establishing that the biosensor behaves like endogenous LC3B in cells, we sought to assess its capacity to dynamically report on LC3B processing. To this end, live cells expressing the WT or the uncleavable G120A biosensor were analyzed using FRET/FLIM. We compared the donor (Aquamarine) lifetime differences between the donor-only and the biosensor to detect FRET events, which are highlighted when the donor lifetime decreases. We then used ΔLifetime as a readout for FRET/FLIM analyses, which was determined by calculating the net difference between the lifetime of the donor-only construct (Aquamarine-proLC3B) and that of a biosensor (either Aquamarine-proLC3B- or proLC3B^G120A^-tdLanYFP). We hypothesized that a positive ΔLifetime would be indicative of a FRET event between Aquamarine and tdLanYFP, therefore corresponding to the presence of unprimed proLC3B.

First, we measured the FRET/FLIM readout of the WT biosensor by calculating its mean ΔLifetime in the total cell area. This includes the cytosol and the puncta structures, in which the precursor, primed and lipidated forms of LC3B are expected to be present. We observed that no FRET was occurring in control cells, as illustrated by a ΔLifetime difference close to zero (Fig. 2A, B). This indicates that the LC3B biosensor is completely primed under basal conditions, leading to the loss of the tdLanYFP moiety. Conversely, the uncleavable G120A biosensor reported a significant increase of ~500 psec in ΔLifetime compared to the WT biosensor. This led us to conclude that the FRET readout of the LC3B biosensor is directly linked to its cleavage on G120. In addition, the FRET readout is specific to the biosensor constructs, as we observed no difference in ΔLifetime between the WT and G120A donor-only constructs (Fig. S2A, B).

**Figure 2.**
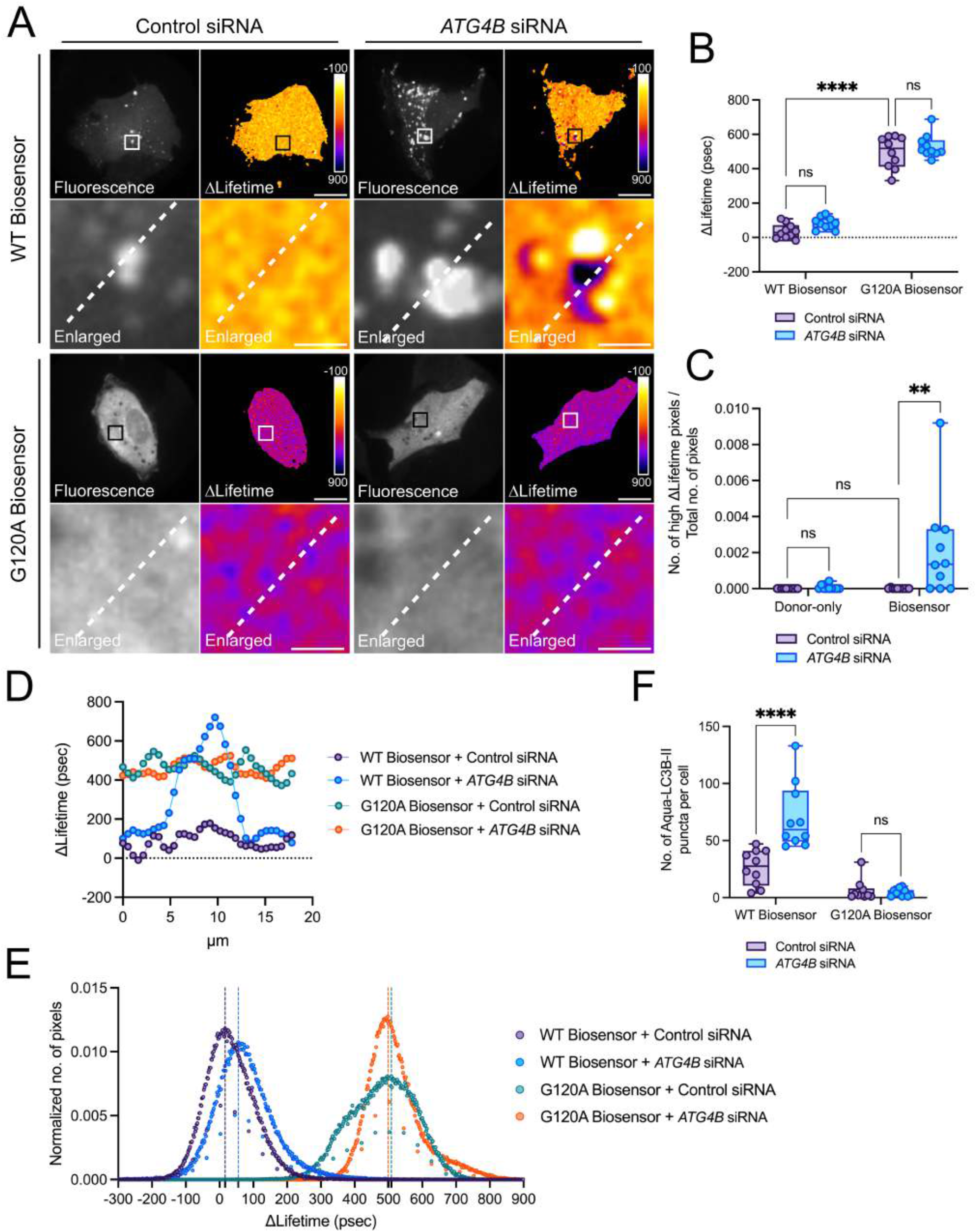
The knockdown of *ATG4B* lowers the priming of the LC3B biosensor. (**A**) Representative fluorescence and ΔLifetime images of U2OS cells co-expressing the WT or G120A biosensor with control or *ATG4B*-specific siRNAs, and analyzed by FRET/FLIM. Squares on the top images of WT or G120A biosensor panels illustrate the location of the enlarged images. Dotted lines on the enlarged images illustrate where the line analysis was performed. Pseudocolor scale: pixel-by-pixel ΔLifetime. Scale bars: overviews, 40 μm; enlarged, 6 μm. Mean ΔLifetime (**B**), number of high ΔLifetime pixels (**C**), line (**D**), histogram (**E**), and number of Aqua-LC3B-II puncta (**F**) analyses of U2OS cells coexpressing the WT or G120A biosensor with control or *ATG4B*-specific siRNAs in (**B**), (**D**), (**E**) and (**F**), and the WT donor or biosensor with control or *ATG4B* siRNA in (**C**). Vertical dotted lines on each histogram depicts the mode value in (**E**). *n* = 10 cells per condition from one representative experiment (of three) in (**B**), (**C**), (**E**) and (**F**). \*\**P* < 0.01, \*\*\*\**P* < 0.0001, ns (not significant) as determined by two-way ANOVA with Tukey’s multiple comparison test in (**B**), (**C**) and (**F**).

We then reasoned that the direct correlation between FRET and priming should make the LC3B biosensor responsive to the presence or absence of ATG4. To this end, we used siRNA-mediated gene silencing to downregulate the *ATG4B* isoform, as it exhibits the highest catalytic efficiency towards LC3B compared to the other members of the ATG4 family [42]. First, we verified the efficiency of the siRNA-mediated downregulation strategy by western blotting, in cells expressing the WT or the G120A biosensor (Fig. S2C, D). When comparing the mean ΔLifetime values, no difference was observed in cells expressing the G120A biosensor under any condition, as expected (Fig. 2A, B). Although no difference in mean ΔLifetime was observable in control or *ATG4B-depleted* cells expressing the WT biosensor (Fig. 2A, B), we noticed the presence of a significant subset of pixels exhibiting high ΔLifetime values only in *ATG4B*-depleted cells (Fig 2A, enlarged ΔLifetime image of the WT biosensor with *ATG4B* siRNA). Therefore, we hypothesized that these pixels might correspond to unprimed proLC3B. To verify our hypothesis, we ascertained that these pixels could be retrieved only upon *ATG4B* downregulation, and that they could represent unprimed pools of LC3B by showing a G120A-like FRET. To this end, we performed a pixel-by-pixel FRET/FLIM analysis to quantify the number of pixels showing ΔLifetime values similar or higher than the mean ΔLifetime of the G120A biosensor. Indeed, *ATG4B* silencing caused a significant increase in the number of pixels with high ΔLifetime values compared to the control condition (Fig. 2C). We then used the power of FRET microscopy to visualize these high ΔLifetime pixels with a line analysis. This analysis allows to observe the local variations in ΔLifetime occurring in the pixels crossed by a straight line. ΔLifetime values along the line went from zero to the levels of the G120A biosensor only in cells silenced for *ATG4B* (Fig. 2A, D). This indicates a lack of proLC3B cleavage occurring locally, and it substantiates the role of ATG4B in this process. We also noticed that the increase in ΔLifetime was localized in pixels found on or near the puncta-shaped structures (Fig. 2A). This reveals the spatial heterogeneity of the priming activity in these areas and uncovers a possible spatial regulation of proLC3B priming, which may be taking place in discrete regions in the vicinity of autophagosomes. When we performed the same analysis for donor-only constructs, we could not detect any high ΔLifetime pixels, ensuring that the effect that we observe with the WT biosensor upon *ATG4B* downregulation was not due to an intrinsic change in the lifetime properties of Aquamarine (Fig. S2E). To make sure that the heterogeneity of ΔLifetime pixels was correctly estimated in the different conditions, we analyzed the data to visualize the ΔLifetime distribution. We hypothesized that the presence of high ΔLifetime pixels with G120A biosensor-like ΔLifetime values should change the overall ΔLifetime distribution. To this end, we superimposed the histograms of the cells expressing the WT biosensor with or without *ATG4B* depletion, and we observed a shift in the histogram mode values of *ATG4B*-depleted cells towards higher ΔLifetime values (Fig. 2E). Since the mode value of a histogram corresponds to the value with the highest frequency, a positive shift in the mode indicates that the ΔLifetime distribution changes due to the presence of FRET events in the biosensor.

We then sought to verify whether FRET events in *ATG4B*-depleted cells were specific to the presence of proLC3B pools in discrete locations, or whether they were due to the clustering of the cleaved reporter. We reasoned that if FRET is due to the unspecific proximity of donor and acceptor molecules, high-ΔLifetime pixels should also be visible when the donor and the acceptor are expressed on distinct LC3B molecules. To rule out this possibility, we compared the FRET behavior of the cells expressing the biosensor with the cells co-expressing the donor and the acceptor constructs [donor-only (Aquamarine-proLC3B) + acceptor-only (proLC3B-tdLanYFP)]. Similar to the WT biosensor, the mean ΔLifetime values of cells co-expressing donor + acceptor in the presence of a control siRNA were close to zero, and no further change was observed upon *ATG4B* downregulation (Fig. S3A-B). Sensitive analyses revealed that donor + acceptor co-expressing cells did not exhibit any high-ΔLifetime pixels when *ATG4B* was depleted, while such high-ΔLifetime pixels were detectable in cells expressing the biosensor (Fig. S3C). As shown in Fig. 2D, *ATG4B* depletion induced a local increase in ΔLifetime values in the proximity of puncta in cells expressing the biosensor (Fig. S3D), and an increase in histogram mode value (Fig. S3E). However, these changes were absent in donor + acceptor co-expressing cells (Fig. S3D, E). These drastic differences in the FRET response of the cells co-expressing the donor and the acceptor compared to the LC3B biosensor strongly suggest that FRET events are intramolecular and specific to the biosensor construct, and they are intimately linked to its priming by ATG4B.

Considering that, in addition to its priming activity, ATG4B also acts as a deconjugating enzyme that governs the ATG8ylation levels [25,43], we checked if its depletion is causing an increase in the puncta-shaped structures. In cells expressing the WT biosensor or donor, we observed a robust increase in the number of LC3B puncta upon *ATG4B* downregulation compared to controls (Fig. 2F, S2F). This indicates that the formation of puncta-shaped structures depends on the presence of ATG4B. As expected, no puncta were observed in cells expressing the G120A biosensor or donor, regardless of the presence or absence of ATG4B (Fig. 2F, S2F). Similarly, when we analyzed protein levels by western blotting, we detected a significant increase in the lipidated levels of the biosensor (Aqua-LC3B-II) and of the endogenous LC3B-II upon *ATG4B* downregulation, compared to the controls (Fig. S2C, D).

Taken together, these results show that the LC3B biosensor allows to visualize changes in the ATG4B-dependent priming of proLC3B. We provide evidence that the biosensor can form LC3B puncta in an ATG4B-dependent manner, demonstrating that our probe recapitulates the key features of endogenous LC3B. Last, we demonstrate that analyzing the FRET response of the biosensor with different modalities allows to monitor the cleavage of proLC3B both at the cellular and at the subcellular level. Our findings support the pertinence of this tool to spatiotemporally characterize LC3B processing in cells.

### The total depletion of ATG4B maximizes the FRET response of the LC3B biosensor

To deepen our understanding of the mode of action of the LC3B biosensor in cells, we used *ATG4* knockout (KO) HeLa cells generated by Agrotis *et al*. using CRISPR/Cas9-mediated approaches [24]. First, we measured the FRET response of the WT or G120A biosensor expressed in control cells. Similarly to what observed in U2OS cells (Fig. 2A, B), the WT biosensor displayed a ΔLifetime close to zero, while the FLIM readout of the cleavage-deficient G120A biosensor showed a ~400 psec ΔLifetime (Fig. 3A, B). When looking at the distribution of the two sensors, we observed that the WT biosensor was capable of forming puncta-like structures while the G120A biosensor showed a cytosolic distribution as expected (Fig. 3A). Overall, the FRET behavior and the distribution of the WT and G120A LC3B biosensors were consistent with our previous observations in U2OS cells.

**Figure 3.**
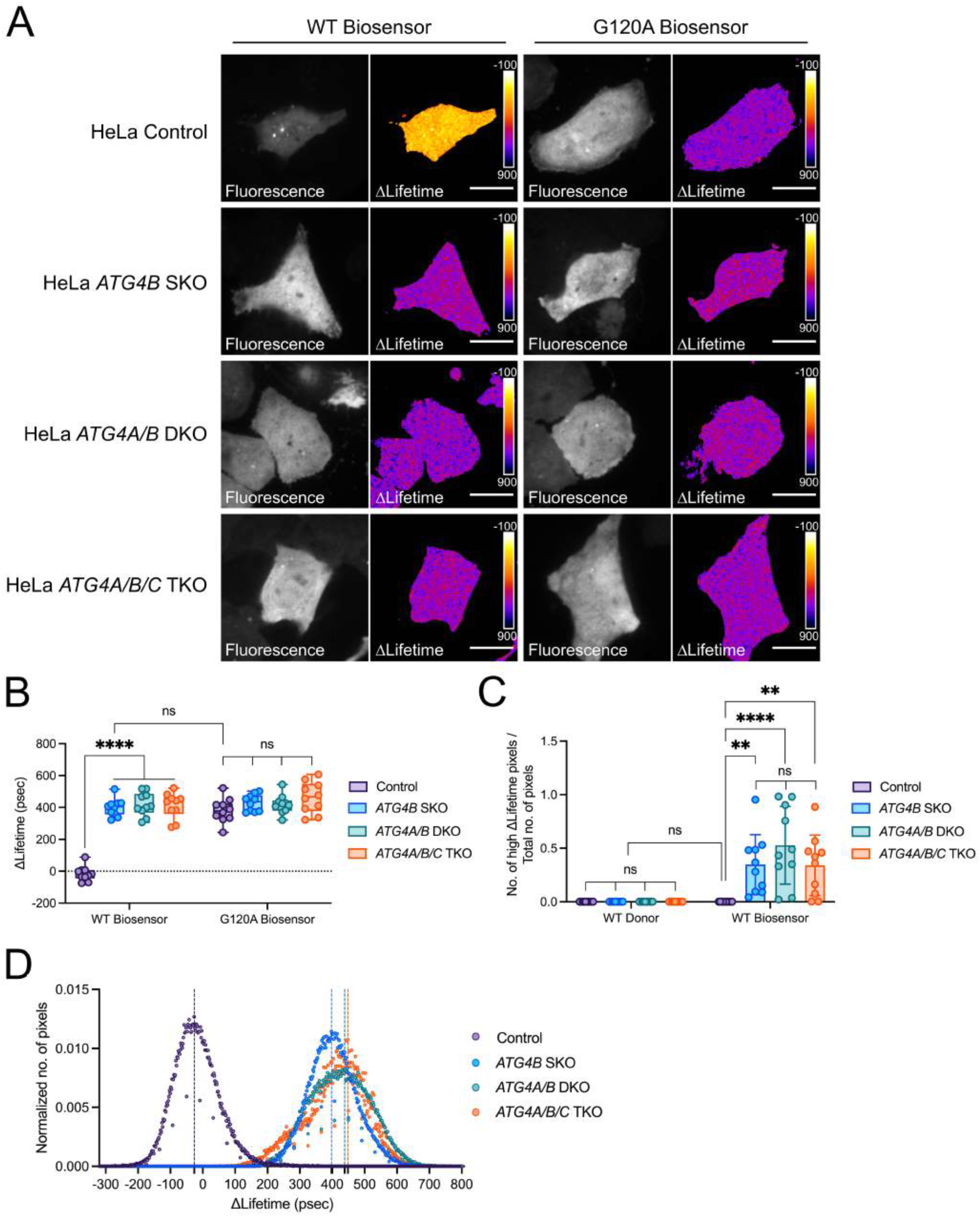
The absence of ATG4B maximizes the FRET response of the LC3B biosensor. (**A**) Representative fluorescence and ΔLifetime images of HeLa control, *ATG4B* SKO, *ATG4A/B* DKO, *ATG4A/B/C* TKO cells expressing the WT or G120A biosensor and analyzed by FRET/FLIM. Pseudocolor scale: pixel-by-pixel ΔLifetime. Scale bar: 40 μm. Mean ΔLifetime (**B**), number of high ΔLifetime pixels (**C**) and histogram (**D**) analyses of HeLa control, *ATG4B* SKO, *ATG4A/B* DKO, *ATG4A/B/C* TKO cells expressing the WT or G120A biosensor. The vertical dotted lines on each histogram depict the mode value in (**D**). *n* = 10 cells per condition from one representative experiment (of three) in (**B**), (**C**) and (**D**). \*\**P* < 0.01, \*\*\*\**P* < 0.0001, ns (not significant) as determined by two-way ANOVA with Tukey’s multiple comparison test in (**B**) and (**C**).

We then sought to explore the FRET/FLIM readout of the LC3B biosensor in cells completely devoid of the ATG4 protease. We expressed the WT biosensor in *ATG4B* single knockout (SKO) HeLa cells and in comparison to control cells, the WT biosensor displayed mean ΔLifetime values of ~400 psec under these conditions (Fig. 3A, B). These values were nearly identical to the mean ΔLifetime of the G120A biosensor. Similarly to the distribution of the G120A biosensor, the WT biosensor in SKO cells did not exhibit noticeable puncta-like structures, and it remained cytosolic. Therefore, these findings support the loss of priming of the WT biosensor in an *ATG4B* SKO background. The lack of priming of the LC3B biosensor was also evident in western blotting analyses (Fig. S4). The WT biosensor expressed in control cells displayed two bands corresponding to the primed and the lipidated forms of the probe, both in the ~43-45 kDa range. In contrast, the WT biosensor expressed in *ATG4B* SKO cells exhibited a single band at ~95 kDa. This band is similar to that observed in cells expressing G120A biosensor, therefore reinforcing the conclusion that the WT biosensor remains unprimed in cells lacking the ATG4B protease. We observed that the complete knockout of *ATG4B* also abolishes the priming of endogenous LC3B, which exhibits a single band at 15 kDa compatible with unprimed proLC3B [44] (Fig. S4). These results indicate that ATG4B is indispensable for the priming and the lipidation of the LC3B biosensor.

We then verified whether the A and C isoforms of ATG4 family could further contribute to the priming of the LC3B biosensor. The mean ΔLifetime values (Fig. 3A, B) and the western blotting profiles (Fig. S4) of the WT biosensor expressed in *ATG4A/B* double knockout (DKO) or *ATG4A/B/C* triple knockout (TKO) HeLa cells [24] showed no differences compared to the *ATG4B* SKO condition. This substantiates previous *in vitro* reports showing that the catalytic activity of ATG4B is maximized towards LC3B [45]. To rule out every possibility that the A or C isoforms could still contribute to the priming of LC3B to some extent, we increased the sensitivity of our analyses by performing pixel-by-pixel FRET/FLIM calculations. We quantified the number of pixels with ΔLifetime values similar or higher than the ΔLifetime of the G120A biosensor in control, *ATG4B* SKO, *ATG4A/B* DKO or *ATG4A/B/C* TKO cells. With this, we aimed to explore subtle changes possibly occurring between the KO cells that may remain undetected in mean ΔLifetime analyses. As expected, we observed a significant increase in the number of pixels with high ΔLifetime in *ATG4B* SKO cells expressing the LC3B biosensor when compared to control cells, further corroborating the results obtained with mean ΔLifetime analyses (Fig. 3C). However, these analyses did not highlight any significant increase in the number of pixels with high ΔLifetime upon further loss of *ATG4A* or *ATG4C* (Fig. 3C) although we noted that *ATG4A/B* DKO and *ATG4A/B/C* TKO cells showed a slight shift towards higher ΔLifetime values in their respective histogram mode value compared to the mode value of *ATG4B* SKO cells (Fig. 3D).

Similarly to what observed for the endogenous LC3B protein, our results demonstrate that the priming of the LC3B biosensor is highly dependent on the presence of ATG4B. When *ATG4B* is absent, the LC3B biosensor displays a significant FRET response, it remains unprimed and cannot be lipidated to form puncta-like structures, overall resembling the behavior of the cleavage- and lipidation-deficient G120A biosensor.

### The ectopic expression of active ATG4B rescues the priming deficiency of the LC3B biosensor in ATG4B-deficient cells

After demonstrating that the priming of the LC3B biosensor is ATG4B-dependent, we then asked whether its FRET response in ATG4B-deficient cells could be rescued by re-expressing ATG4B. To this end, we co-expressed the LC3B biosensor together with an empty vector or with a vector coding for WT ATG4B. Then, we evaluated the FRET/FLIM behavior of these conditions both in control and in *ATG4B* SKO cells. Consistent with our previous findings (Fig. 3), the LC3B biosensor co-expressed with an empty vector in *ATG4B* SKO cells showed a significant increase in its mean ΔLifetime values compared to control cells (Fig. 4A, B). Conversely, the expression of WT ATG4B in *ATG4B* SKO cells caused a drastic decrease in the mean ΔLifetime values of the LC3B biosensor, which were close to zero. This suggests that the reintroduction of WT ATG4B is sufficient to fully rescue the cleavage of the LC3 biosensor in a SKO background. Upon the expression of exogenous ATG4B in control cells, we also observed that the distribution of the biosensor was cytosolic and without significant puncta-like structures (Fig. 4A). On the contrary, control cells co-expressing an empty vector were capable of forming puncta-like structures. This is in agreement with previous reports showing that the overexpression of exogenous ATG4B blocks the lipidation of LC3B and by doing so, it leads to the disappearance of LC3-positive puncta in cells [13,46,47].

**Figure 4.**
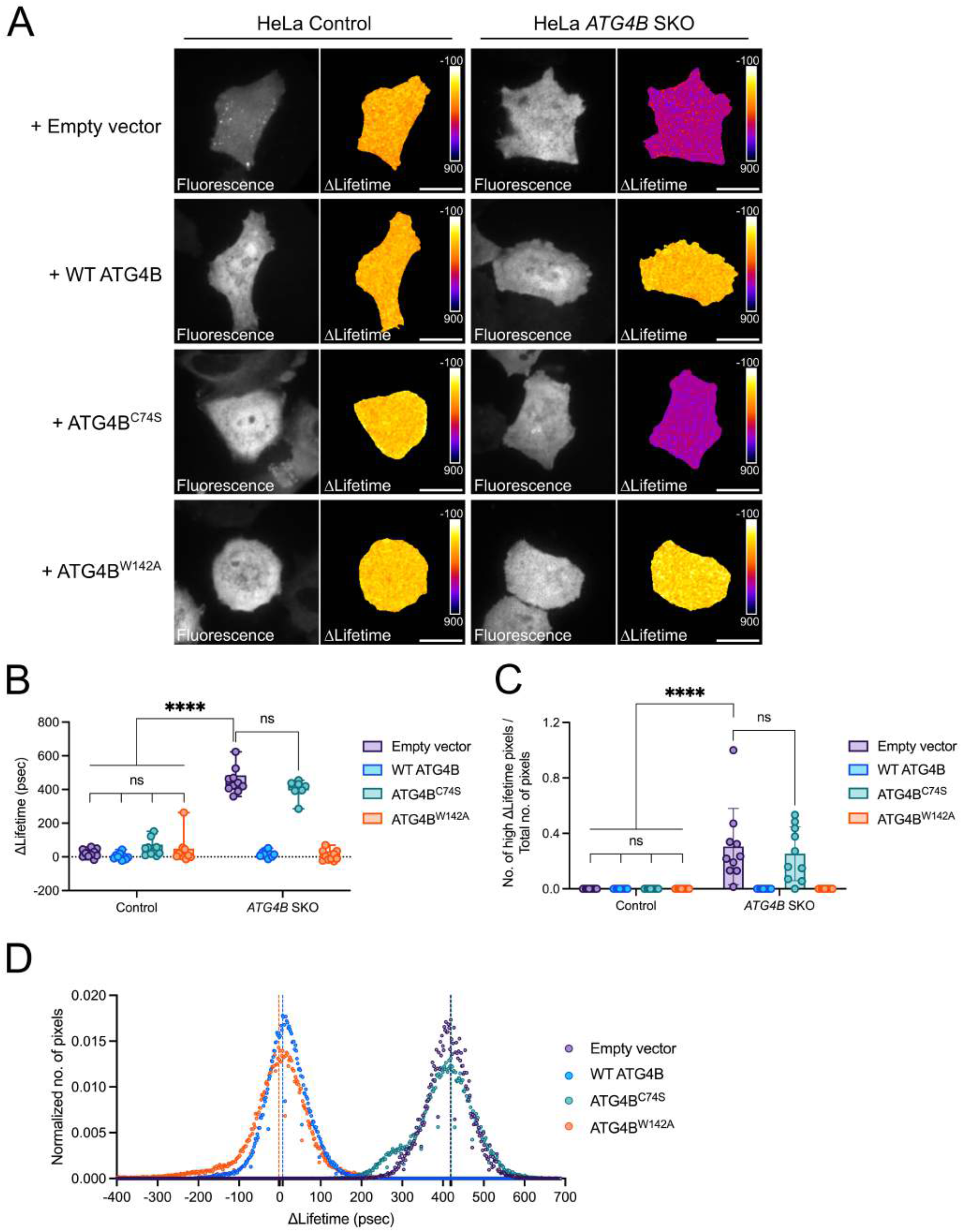
The priming deficiency of the LC3B biosensor is rescued when expressing WT or ATG4B^W142A^ in *ATG4B* SKO cells. (**A**) Representative fluorescence and ΔLifetime images of control and *ATG4B* SKO HeLa cells co-expressing the WT biosensor with an empty vector, or with vectors expressing WT ATG4B, ATG4B^C74S^ or ATG4B^W142A^, and analyzed by FRET/FLIM. Pseudocolor scale: pixel-by-pixel ΔLifetime. Scale bar: 40 μm. Mean ΔLifetime (**B**) and number of high ΔLifetime pixels (**C**) analyses of control and *ATG4B* SKO cells co-expressing the WT biosensor with an empty vector, or with vectors expressing WT ATG4B, ATG4B^C74S^ or ATG4B^W142A^. (**D**) The histogram analysis of *ATG4B* SKO cells coexpressing the WT biosensor with an empty vector, or with vectors expressing WT ATG4B, ATG4B^C74S^ or ATG4B^W142A^. Vertical dotted lines on each histogram depict the mode value in (**D**). *n* = 10 cells per condition from one representative experiment (of three) in (**B**), (**C**) and (**D**). \*\*\*\**P* < 0.0001, ns (not significant) as determined by two-way ANOVA with Tukey’s multiple comparison test in (**B**) and (**C**).

Next, we assessed whether two mutant forms of ATG4B with different catalytic activities could rescue the priming deficiency of the LC3B biosensor in *ATG4B* SKO cells. We first explored the consequences of mutating the Cys74 residue of ATG4B into Ser (C74S). Cys74 belongs to a group of three aminoacids known as the “catalytic triad”, and its mutation into Ala or Ser was shown to cause a complete loss of the catalytic activity of ATG4B [13,35]. When ATG4B^C74S^ was co-expressed with the LC3B biosensor in *ATG4B* SKO cells, we measured a mean ΔLifetime comparable to *ATG4B* SKO cells transfected with an empty vector (Fig. 4A, B). This indicates that the catalytically-dead C74S mutant was unable to cleave and, by consequence prime, the LC3B biosensor. We then tested a second mutant form of ATG4B where Trp142 was mutated into Ala (W142A). Trp142 is one of the residues surrounding the catalytic triad, and its mutation into Ala was reported to significantly reduce the catalytic activity of ATG4B *in vitro* [35]. Surprisingly, we observed that ATG4B^W142A^ behaved similarly to WT ATG4B when expressed in SKO cells, and it resulted in a complete cleavage of the LC3B biosensor in FRET/FLIM analyses (Fig. 4A, B). This supports the superior catalytic efficiency of ATG4B to prime proLC3B in living cells, even in conditions where its catalytic activity is severely compromised by the W142A mutation. To verify that these changes in mean ΔLifetime were specific to the LC3B biosensor, we analyzed the mean ΔLifetime profiles of control or *ATG4B* SKO cells expressing the donor-only Aquamarine-LC3B construct together with WT, C74S or W142A ATG4B (Fig. S5A). As expected, we did not observe any difference among all the conditions tested, further confirming that the mean ΔLifetime FRET/FLIM readout is specific towards the ATG4B-mediated cleavage of the LC3B biosensor.

We then wanted to verify if our approach based on mean ΔLifetime was sensitive enough to conclude on the capacity of the W142A mutant to fully prime LC3B. Therefore, we further evaluated the readout of the LC3B biosensor by performing pixel-by-pixel FRET/FLIM calculations, which we previously showed to be more sensitive than mean ΔLifetime analysis (Fig. 2). As expected, we observed a significant increase in the number of pixels with high ΔLifetime – indicating no LC3B priming – in *ATG4B* SKO cells co-expressing an empty vector or ATG4B^C74S^ compared to control cells (Fig. 4C). Similarly to mean ΔLifetime analyses (Fig. 4B), expressing WT ATG4B or ATG4B^W412A^ in *ATG4B* SKO cells revealed an absence of pixels with high ΔLifetime values, suggesting a complete rescue of the priming activity of the LC3B biosensor with both constructs (Fig. 4C). Accordingly, the histogram analyses of the distribution of FRET pixels in all conditions showed almost identical mode values in *ATG4B* SKO cells co-expressing WT or ATG4B^W142A^, with their respective mode values centered around zero (Fig. 4D). On the contrary, the mode values of *ATG4B* SKO cells expressing an empty vector or ATG4B^C74S^ were drastically shifted towards 400 psec, which is again indicative of significant FRET (Fig. 4D). As expected, the analysis of the high ΔLifetime pixels in control or SKO cells expressing the donor-only Aquamarine-LC3B and each of the ATG4B constructs did not reveal any difference (Fig. S5B). Again, this substantiates the specificity of our different FRET/FLIM readouts for the LC3B biosensor only. In addition to this, the WT biosensor expressed in control cells with each of the ATG4B constructs exhibit similar histogram mode values (Fig. S5C). Nevertheless, we observed the presence of a shoulder corresponding to ΔLifetime values of ~200-300 psec in control cells co-expressing the WT biosensor with ATG4B^C74S^ (Fig. S5C), which could reflect the previously reported dominant negative effects of ATG4B^C74S^ on the soluble forms of LC3B [46].

Finally, to rule out if the capacity of WT or mutant ATG4Bs to prime proLC3B can differ due to changes in their expression levels, we performed western blotting analyses to compare exogenous ATG4B protein levels and their proLC3B priming patterns. We first verified that protein expression levels of overexpressed WT or mutant ATG4Bs in control or *ATG4B* SKO cells were comparable (Fig. S5D). Then, we confirmed that *ATG4B* SKO cells co-expressing WT or ATG4B^W142A^ but not ATG4B^C74S^ exhibited bands compatible with the molecular weight of Aquamarine + LC3B-I at ~45 kDa. On the hand, *ATG4B* SKO cells co-expressing ATG4B^C74S^ only exhibited a higher molecular weight band compatible with the size of Aquamarine + proLC3B + tdLanYFP (Fig. S5D).

Altogether, our results demonstrate that the proLC3B priming deficiency observed in *ATG4B* SKO cells can be fully restored when co-expressing the WT or W142A forms of ATG4B, but not with the catalytically-dead C74S. Although ATG4B^W142A^ was shown to display a reduced catalytic activity *in vitro* [35], we demonstrate that this mutant is able to prime proLC3B in living cells similar to WT ATG4B. These data were obtained with three independent methods to calculate the FRET behavior of the LC3B biosensor and with an orthogonal approach based on western blotting analyses. Together, they provide novel insights on the superior capacity of ATG4B to prime LC3B *in cellulo* even in conditions where its catalytic activity is compromised. Importantly, they also support the notion that the catalytic activity of ATG4B needs to be completely eliminated to abolish LC3B priming.

### The LC3B biosensor reveals the mode of action of ATG4 inhibitors in cells

Given that the biosensor reports on the priming of LC3B by ATG4B, we sought to investigate whether it is also capable to respond to pharmacological compounds that inhibit the ATG4B-LC3B axis. We first explored the readout of the LC3B biosensor on commercially-available inhibitors of ATG4s. Tioconazole, LV-320, FMK-9a, NSC 185058 (NSC) and *Z-L-Phe* chloromethyl ketone (ZPCK) were evaluated in their capacity to inhibit the priming and/or deconjugation activities of ATG4B [48–53]. These inhibitors have either been synthesized or identified in screening studies, and they were previously shown to inhibit ATG4B and/or other ATG4 isoforms. They were also reported to have a significant therapeutic potential in chemotherapy-resistant cancer subtypes with elevated levels of autophagy [29,30].

We determined the working concentration of LV-320, FMK-9a and NSC based on previous reports testing the effects of these compounds in cells [49,51,52]. For Tioconazole, we decreased the concentration (to 4 μM) as we experienced high rates of cell death when using the compound at the previously reported concentration (40 μM) [48]. Finally, for ZPCK, we determined a dose that can be tolerated by our cells and based on available IC50 values determined by different assays [53]. Since the reported concentrations are different according to the compound used, we then chose to standardize the duration of the treatment to 6 hours to be able to detect short- to mid-term effects of each compound. After determining the dose and the duration to be tested, we first measured the mean ΔLifetime values in HeLa cells expressing the WT or the G120A biosensor, and treated or not with each of the five inhibitors. When cells are treated with the inhibitors, we expected to observe a FRET response compatible to that detected in cells downregulated for *ATG4B* as the available reports demonstrated the effects of ATG4 inhibitors mostly inhibiting the deconjugation activity with limited impact on the priming activity [48,49,51,52],. Within our selected set of inhibitors, two of them – NSC and ZPCK – were found to significantly increase mean ΔLifetime values compared to controls (Fig. 5A vs. 5E, F). On the contrary, Tioconazole, LV-320 and FMK-9a did not alter the mean ΔLifetime values of the biosensor (Fig. 5A-D). However, the mean ΔLifetime values of cells treated with NSC or ZPCK remained lower than those of cells expressing the G120A biosensor. This suggests that the inhibitory effect of these drugs towards ATG4B remains partial. We then asked whether the main function of NSC and ZPCK is to inhibit the ATG4B-dependent deconjugation of LC3B or its priming. We observed a significant increase in the number of Aqua-LC3B-II puncta structures upon treatment with NSC or ZPCK, further supporting the idea that autophagy can still be triggered in the presence of these compounds and that the inhibition of ATG4B priming activity is not complete (Fig. 5E, F). However, increased levels of Aqua-LC3B-II puncta structures were observed in cells treated with Tioconazole, LV-320 or FMK-9a as well (Fig. 5B-D), and we found no correlation between puncta numbers and mean ΔLifetime values with any of the inhibitors (Fig. S6). These findings suggest that all the tested compounds preferentially hinder the deconjugation activity of ATG4B towards LC3B, which was described to be more sensitive to ATG4B inhibition than the priming activity [54]. In addition to deconjugation, mean ΔLifetime differences provide the first evidence that NSC and ZPCK also inhibit the priming activity of ATG4B.

**Figure 5.**
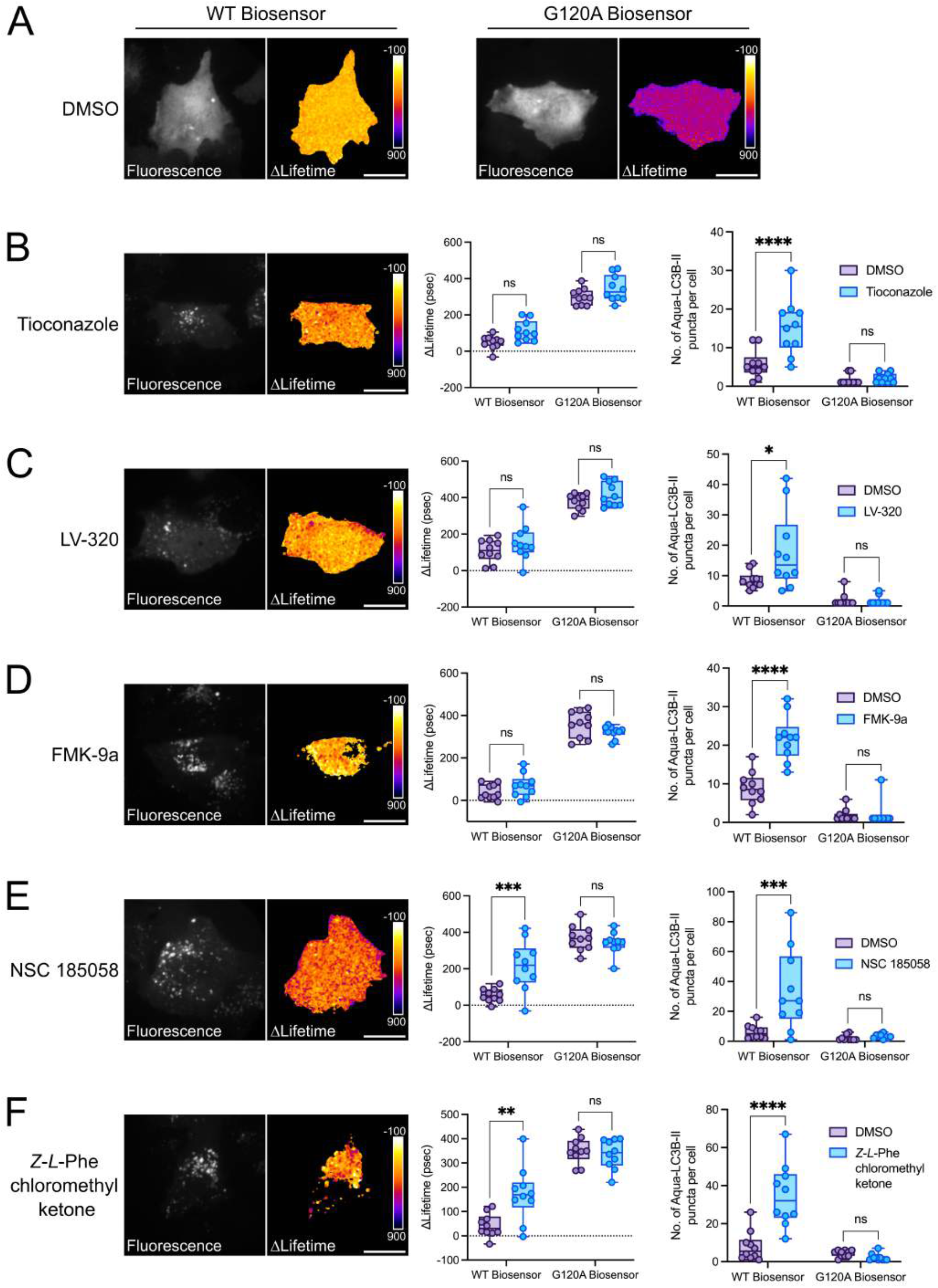
ATG4B inhibitors variably alter the ΔLifetime behavior and LC3B puncta number in cells expressing the LC3B biosensor. (**A**) Representative fluorescence and ΔLifetime images of HeLa cells expressing the WT or G120A biosensor, treated with DMSO (6h), and analyzed by FRET/FLIM. Representative fluorescence and ΔLifetime images of HeLa cells expressing the WT biosensor and treated with Tioconazole (6h, 4 μM) (**B**), LV-320 (6h, 120 μM) (**C**), FMK-9a (6h, 10 μM) (**D**), NSC 185058 (6h, 100 μM) (**E**), or *Z*-*L*-Phe chloromethyl ketone (6h, 3 μM) (**F**). Mean ΔLifetime and number of Aqua-LC3B-II puncta analyses of HeLa cells expressing the WT or G120A biosensor and treated with Tioconazole (6h, 4μM) (**B**), LV-320 (6h, 120 μM) (**C**), FMK-9a (6h, 10 μM) (**D**), NSC 185058 (6h, 100 μM) (**E**), or *Z*-*L*-Phe chloromethyl ketone (6h, 3 μM) (**F**). Pseudocolor scale: pixel-by-pixel ΔLifetime. Scale bars: 40 μm. *n* = 10 cells per condition from one representative experiment (of three) in (**B**-**F**). \**P* < 0.05, \*\**P* < 0.01, \*\*\**P* < 0.001, \*\*\*\**P* < 0.0001, ns (not significant) as determined by two-way ANOVA with Tukey’s multiple comparison test in (**B-F**).

To dissect the efficacy of the five ATG4B inhibitors with methods allowing for greater sensitivity than mean ΔLifetime, we then performed high ΔLifetime-pixel counting and histogram analyses. The high ΔLifetime-pixel counting analyses revealed that not only NSC and ZPCK, but also FMK-9a exhibited a significantly increased number of pixels with high ΔLifetime compared to control cells (Fig. 6A-D). We also performed western blotting analyses to investigate if proLC3B accumulation can be observed upon treatment with ATG4B inhibitors. In line with FRET results, we observed bands at a molecular weight compatible with the unprimed biosensor (Aqumarine + proLC3B +tdLanYFP) only in cells treated with FMK-9a, NSC or ZPCK but not in cells treated with Tioconazole or LV-320 (Fig. S7). However, we also noticed that the proLC3B band is more abundant in cells treated with FMK-9a or NSC compared to cells incubated with ZPCK. Although inhibitors were added in all steps of sample preparation to avoid proLC3B priming at any moment, it is likely that the differences observed in the abundance of proLC3B bands is due to the differential efficacy of the compounds used under denaturing conditions. Therefore, these results further substantiate the importance of quantitative assays performed with living cells when investigating the efficacy of potential ATG4B inhibitors. Indeed, when we analyzed the distribution of ΔLifetime pixels on histogram analyses, we observed that approximately 10-20% of pixels in cells treated with NSC or ZPCK were exhibiting G120A biosensor-like ΔLifetime values (Fig. 6A-D). These values were lowered in the presence of Tioconazole, LV-320 and FMK-9a, further corroborating the superiority of NSC and ZPCK in inhibiting ATG4B (Fig. S8). In line with these findings, histogram analyses revealed that NSC (113 psec) and ZPCK (150 psec) have the largest histogram mode value shift from that of control cells (Fig. 6C, D). Interestingly, these analyses showed that FMK-9a, Tioconazole and LV-320 also display a mode value shift from that of control cells, respectively of 74, 65 and 47 psec (Fig. 6B and S8B-C). Therefore, this sensitive analysis method indicates a mild inhibition of the priming activity of ATG4B by FMK-9a, Tioconazole and LV-320 as well, which was undetectable with the other analysis methods. Furthermore, this approach substantiates the superior capacity of NSC and ZPCK in inhibiting ATG4B.

**Figure 6.**
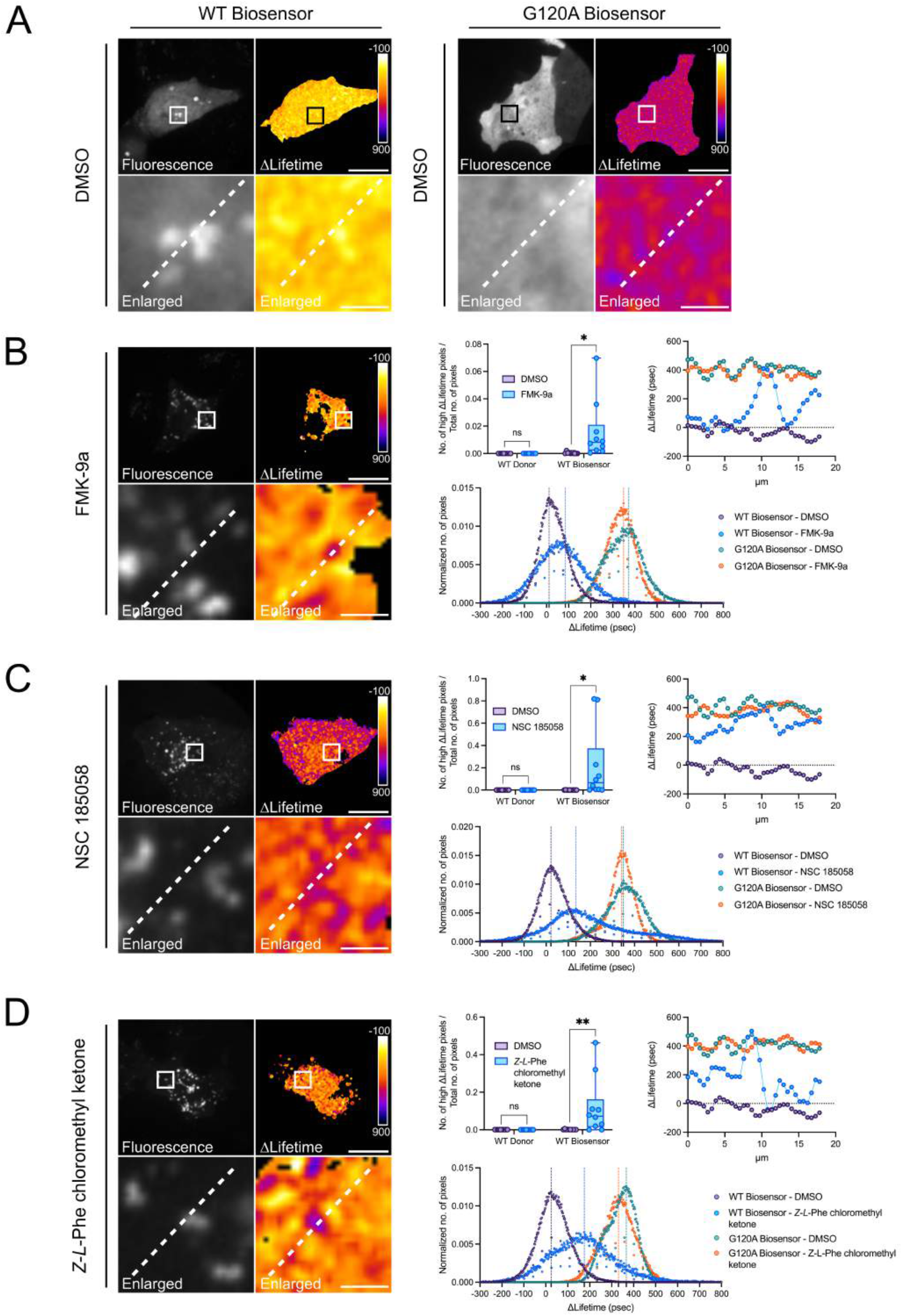
The LC3B biosensor reveals the mode of action of FMK-9a, NSC 185058 and *Z*-*L*-Phe chloromethyl ketone in cells. (**A**) Representative fluorescence and ΔLifetime images of HeLa cells expressing the WT or G120A biosensor, treated with DMSO (6h), and analyzed by FRET/FLIM. Representative fluorescence and ΔLifetime images of HeLa cells expressing the WT biosensor and treated with the following compounds: FMK-9a (6h, 10 μM) (**B**), NSC 185058 (6h, 100 μM) (**C**), or *Z-L-Phe* chloromethyl ketone (6h, 3 μM) (**D**). Squares on the top images of WT or G120A biosensor panels illustrate the location of the enlarged images. Dotted lines on the enlarged images illustrate where the line analysis was performed. Pseudocolor scale: pixel-by-pixel ΔLifetime. Scale bars: overviews, 40 μm; enlarged, 6 μm. Number of high ΔLifetime pixels analysis of HeLa cells expressing the WT donor or biosensor and treated with FMK-9a (6h, 10 μM) (**B**), NSC 185058 (6h, 100 μM) (**C**), or *Z*-*L*-Phe chloromethyl ketone (6h, 3 μM) (**D**). Line and histogram analyses of HeLa cells expressing the WT or G120A biosensor and treated with FMK-9a (6h, 10 μM) (**B**), NSC 185058 (6h, 100 μM) (**C**), or *Z*-*L*-Phe chloromethyl ketone (6h, 3 μM) (**D**). *n* = 10 cells per condition from one representative experiment (of three) in (**B**-**D**). *\*P* < 0.05, *\*\*P* < 0.01, ns (not significant) as determined by two-way ANOVA with Tukey’s multiple comparison test in (**B-D**).

Since we observed the presence of high ΔLifetime pixels with all the inhibitors, we then sought to investigate the subcellular location of these pixels using line analysis. Similarly to what observed on cells silenced for *ATG4B* (Fig. 2), we noticed that high ΔLifetime pixels were located either on the puncta-shaped structures, or in the surrounding area (Fig. 6 and S8). Line analyses also revealed that these pixels had ΔLifetime values comparable to those of the G120A biosensor, regardless of the compound used. Taken together, these findings show that ATG4B inhibition using commercially-available drugs reduces the priming rates of proLC3B at discrete sites, where unprimed LC3B reservoirs can be found within or in the proximity of puncta-shaped structures.

Afterwards, we explored the effect of MG132, a peptide aldehyde that inhibits both the proteasome and cysteine proteases [55,56]. Considering that ATG4B is a cysteine protease [14,35], we reasoned that MG132 might be able to block its catalytic activity towards LC3B. However, its inhibitory capacity towards ATG4B has never been explored. Therefore, we used our FRET pipeline to explore the efficacy of an inhibitor with unknown effects towards ATG4B. Therefore, we treated HeLa cells expressing the WT or G120A biosensor with DMSO or MG132, and calculated their mean ΔLifetime (Fig. S9A, B). FLIM analyses revealed a significant increase in the mean ΔLifetime values of the WT biosensor in the presence of MG132. We reasoned that MG132 could promote FRET within the biosensor either by inhibiting the priming activity of ATG4B, or by inhibiting LC3B degradation by the proteasome. To distinguish between these two possibilities, we relied on the subcellular distribution of the biosensor. In this light, the biosensor should be retrieved in the cytosol in case MG132 had mainly an ATG4B-specific inhibition on proLC3B priming. Alternatively, it should rather be found in puncta-like structures if this drug acted as a proteasome inhibitor. We observed a significant increase in the number of Aqua-LC3B-II puncta-like structures in cells expressing the WT biosensor and treated with MG132, when compared to DMSO-treated cells (Fig. S9B). Therefore, this localization of the sensor in puncta-like structures suggests that MG132 is more efficient as a proteasome inhibitor, rather than a specific ATG4B inhibitor. Although the number of pixels with high ΔLifetime values did not significantly increase upon MG132 treatment (Fig. S9B), the histogram analysis of MG132-treated cells expressing the WT biosensor revealed a mode value change when compared to DMSO-treated cells and to cells expressing the G120A biosensor (Fig. S9C). Overall, this indicates that G120A-like FRET events are quantitatively modest in the presence of MG132, suggesting that MG132 preferentially acts as a proteasomal inhibitor rather than an ATG4B-specific inhibitor. Although G120A-like FRET events were limited under these conditions, we sought to explore their spatial localization. Line analyses performed in cells expressing the WT biosensor and treated with MG132 revealed that the ΔLifetime variations of the biosensor were of ~200 psec in the cytosol, and reaching ΔLifetime values of the G120A biosensor (~400 psec) on or near the LC3B puncta (Fig. S9B, D). In contrast, no fluctuations were observed in the cytosol or near puncta in control cells. These two FRET values observed after treating cells with MG132 could be recapitulative of the dual action of this compound: a modest ATG4B inhibitor keeping the biosensor in the cytosol, and a more potent proteasomal inhibitor on LC3B-positive puncta. Of note, since Aqua-LC3B can efficiently localize on puncta-like structures (Fig. S9B) which display high FRET (Fig. S9B, D), our data raise the possibility that MG132 does not alter the cleavage of tdLanYFP from the biosensor. It is possible that the inhibition of proteasomal activity impairs the degradation of the FRET acceptor, thereby allowing for non-specific FRET events. Indeed, western blotting of cells treated with MG132 exhibited no band that may be compatible with the size of the unprimed biosensor (Fig. S9E). Therefore, these results underline the importance of performing spatially-resolved, pixel-by-pixel FRET calculations to understand where and to what extent the biosensor is active. Last, they also highlight the poor efficacy of MG132 as an ATG4B-specific inhibitor.

Overall, our data demonstrate that the LC3B FRET biosensor is a powerful tool to evaluate the mode of inhibition of ATG4B-specific compounds. We also provide an innovative methodology where individual sets of microscopy data can be analyzed using three independent approaches. Cumulating the information obtained by the three approaches allows to spatially localize and quantify the ATG4B-dependent priming and deconjugation of LC3B with unprecedented precision, and it is mandatory to characterize the mode of action of present and future ATG4B-specific inhibitors.

### The LC3B biosensor uncovers the CDK1-dependent regulation of the ATG4B-LC3B axis at mitosis

Given the sensitivity of our biosensor to monitor autophagy within the ATG4B-LC3B axis, we sought to explore the involvement of this nexus in a paradigm relevant for cell physiology. In this light, we assessed the FRET behavior of the LC3B biosensor at mitosis, a cell cycle phase where the involvement of autophagy is still controversial. Although autophagy was described to be turned off during cell division [57–59], numerous studies reported the presence of LC3B-positive puncta in mitotic cells [60–62], and the role of ATG4B in this cell cycle phase and on those puncta has never been elucidated. As we uncovered the presence of unprimed LC3B pools on or in the close vicinity of puncta-shaped structures upon *ATG4B* downregulation or ATG4B inhibition, we asked whether mitotic LC3B puncta could be discrete sites containing unprimed LC3B. To this end, we first compared the FRET response of the WT biosensor between interphase cells (unsynchronized), and cells arrested at G2/M following treatment with nocodazole and then released to reach metaphase. When compared to cells expressing the G120A biosensor, the mean ΔLifetime values of the WT biosensor both in unsynchronized and in nocodazole-treated cells were close to zero (Fig. 7A-B). Though LC3B puncta-shaped structures were present in mitotic cells as previously reported [60–62], high-ΔLifetime pixel analyses revealed that proLC3B-like pixels were absent in these cells (Fig. 7C). Line analysis, on the other hand, identified local ΔLifetime variations of ~200 psec on or around the puncta of nocodazole-treated mitotic cells (Fig. 7D). These events were quantitatively modest in number, as the histogram mode value of mitotic cells following nocodazole-mediated synchronization fluctuates around zero (Fig. 7E). Altogether, these results suggest that LC3B is mainly cleaved in mitotic cells, and that the mitotic repression of autophagy is not related with the accumulation of proLC3B pools.

**Figure 7.**
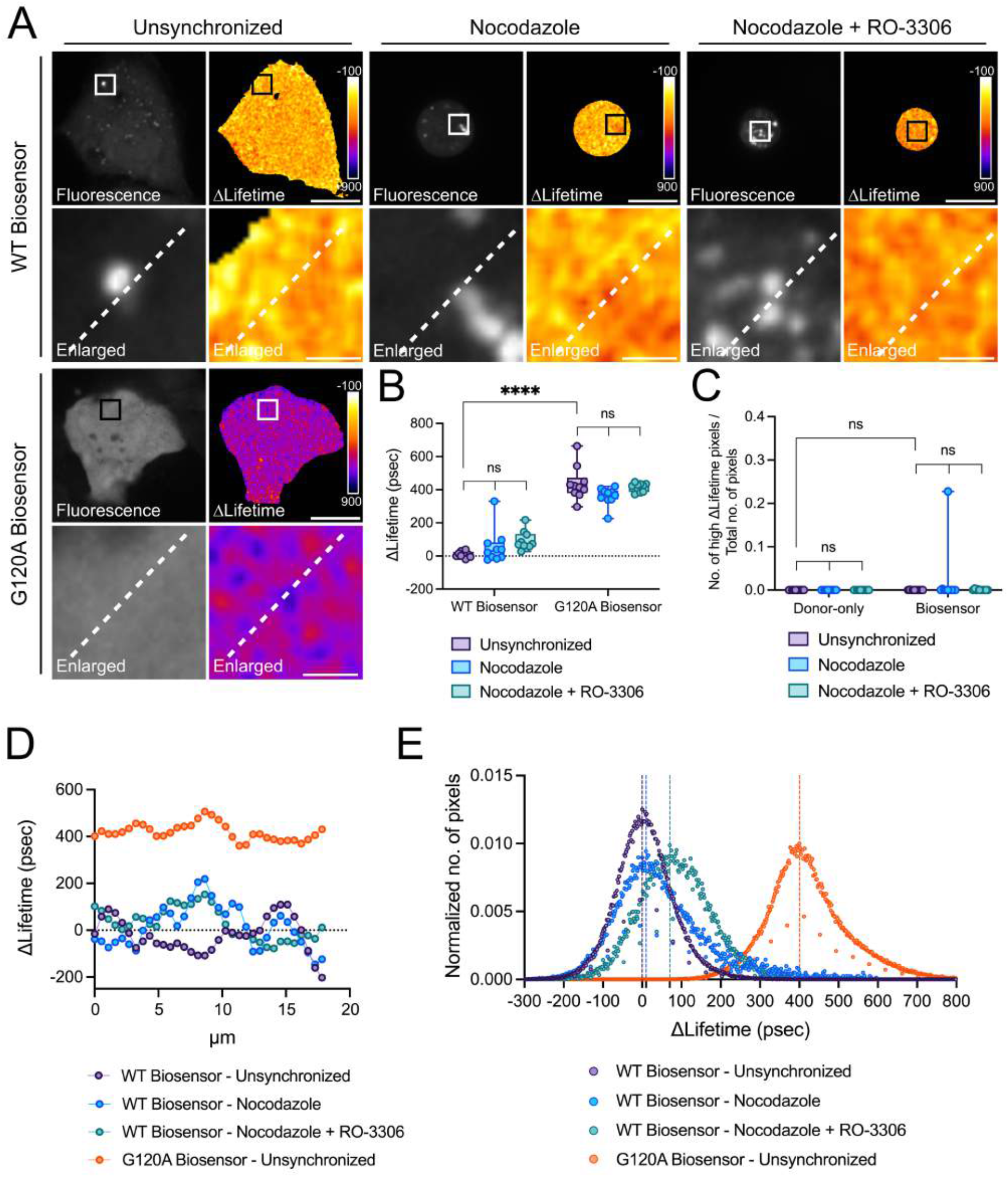
The LC3B biosensor reports on the CDK1-dependent regulation of the ATG4B-LC3B axis at mitosis (**A**) Representative fluorescence and ΔLifetime images of unsynchronized U2OS cells expressing the WT or G120A biosensor, or cells expressing the WT biosensor and treated with nocodazole-only (16h, 100 ng/ml) or co-treated with nocodazole (16h, 100 ng/ml) and RO-3306 (2h, 2 μM), and analyzed by FRET/FLIM. Squares on the top images of WT or G120A biosensor panels illustrate the location of the enlarged images. Dotted lines on the enlarged images illustrate where the line analysis was performed. Pseudocolor scale: pixel-by-pixel ΔLifetime. Scale bars: overviews, 40 μm; enlarged, 6 μm. Mean ΔLifetime (**B**), number of high ΔLifetime pixels (**C**), line (**D**) and histogram (**E**) analyses of unsynchronized, or nocodazole-only (16h, 100 ng/ml), or nocodazole (16h, 100 ng/ml) and RO-3306 (2h, 2 μM) treated U2OS cells expressing the WT or G120A biosensor in (**B**), (**D**) and (**E**), and the WT donor or biosensor in (**C**). Vertical dotted lines on each histogram depicts the mode value in (**E**). *n* = 10 cells per condition from one representative experiment (of three) in (**B**), (**C**) and (**E**). *\*\*\*\*P* < 0.0001, ns (not significant) as determined by two-way ANOVA with Tukey’s multiple comparison test in (**B**) and (**C**).

In a recent study published by Odle *et al*., it has been shown that CDK1 ensures the mitotic repression of autophagy by replacing mTORC1-dependent inhibitory phosphorylations on autophagy regulator proteins such as ATG13, ULK1, ATG14 and TFEB [63]. This report also showed that the treatment of mitotic cells with the CDK1 inhibitor RO-3306 reversed the inhibitory phosphorylations on autophagy regulator proteins. However, it is currently unknown whether CDK1 inhibition also plays a role within the ATG4B-LC3B axis at mitosis, potentially by regulating the ATG4B-dependent proLC3B processing. To this end, we synchronized U2OS cells at mitosis with nocodazole and we inhibited CDK1 with RO-3306. First, we confirmed the efficacy of CDK1 inhibition, using the activation of the cell cycle protein AURKA as a readout. Indeed, AURKA was shown to be activated on Thr288 at mitosis in a CDK1-dependent manner [64]. Therefore, we used our FRET biosensor reporting on AURKA activation [39] to explore the effect of RO-3066 at mitosis. With this tool, we observed that the mitotic spindle failed to form correctly in U2OS cells synchronized at mitosis and treated with RO-3066 (Fig. S10A). This is comparable to what observed with a kinase-dead mutant of the AURKA biosensor in previous reports [38,39], indicating mitotic defects compatible with a failure in activating AURKA [64]. As expected, the FRET readout of the AURKA biosensor revealed a lowered activation of AURKA in cells treated with RO-3066 (Fig. S10A-B). After confirming that CDK1 is inhibited under our experimental conditions, we explored the FRET readout of the LC3B biosensor in cells synchronized at mitosis and in the presence or absence of RO-3306. Although we noticed a trend towards higher mean ΔLifetime values in nocodazole and RO-3306 co-treated cells as compared to either unsynchronized or nocodazole-only treated cells, this trend was not significant (Fig. 7A-B). Similarly, the sensitive high-ΔLifetime pixel analyses did not reveal any significant changes in nocodazole and RO-3306 co-treated cells when compared to unsynchronized or nocodazole-only treated cells (Fig. 7C). However, line analyses showed ~100/200 psec local ΔLifetime variations (Fig. 7D). These differences were abundant in number, since the histogram mode value of nocodazole and RO-3306 co-treated cells shifted towards high ΔLifetime values (Fig. 7E). This increase suggests a CDK1-dependent stalling of the proLC3B processing rates by ATG4B. Overall, these results uncover the role of CDK1 in regulating the ATG4B-LC3B axis at mitosis. In addition, they further highlight the superior sensitivity of the LC3B biosensor to explore subtle changes in the regulation of autophagy, and particularly in paradigms where the role of the ATG4B-LC3B axis still remains to be determined.

## Discussion

In this study, we demonstrated that the LC3B biosensor is a robust tool to monitor autophagy, as it responds to the priming and deconjugation activities of ATG4B on LC3B. We showed that these functions of ATG4B can be followed by using one single probe with a dual readout based on FRET, and on the accumulation of the probe on autophagosomes.

Among the several approaches available to monitor autophagy, the most widely used assays rely on the use of single fluorescent protein (FP)-tagged LC3B probes to quantify the number of autophagosomes [65]. The LC3B biosensor retains this property, since it functions as a standard single FP-tagged probe after LC3B is primed. We also show that the biosensor reports on autophagy induction and/or inhibition while colocalizing with the lysosomal marker protein LAMP2 in an autophagy-dependent manner, similarly to other LC3B-based fluorescent constructs. Since the LC3B biosensor is constituted of a pair of FPs resistant to acidic pH, its readout can be followed throughout the entire autophagy pathway. Importantly, the LC3B biosensor has the capacity to respond to proLC3B priming in living cells, thanks to these FPs behaving as a donor-acceptor FRET pair. The proLC3B priming by ATG4s is among the earliest events occurring when autophagy is triggered [13,36]. A FRET-based strategy relying on a CFP/YFP donor-acceptor pair has already been used to measure the enzymatic activity of ATG4A and ATG4B towards the ATG8 family in a purely *in vitro* system [66]. However, this strategy has never been implemented in living cells, most likely due to the lack of yellow FPs retaining their acceptor properties in conditions of acidic pH. The recent development of tdLanYFP [39] allowed us to create an LC3B biosensor suitable for living cells. By following the FRET behavior of the biosensor, we showed that the probe responds to the ATG4B-dependent changes in proLC3B priming. It was previously reported that proLC3B is primed nearly instantaneously after translation, due to the constitutive proteolytic activity of ATG4B [36,67]. In line with this, we found that the LC3B biosensor was almost completely primed under basal conditions, without any detectable accumulation of proLC3B in cells. In contrast, we showed that the proLC3B priming activity is altered in cells silenced for *ATG4B*, and that the unprimed biosensor is located on or in close vicinity of puncta-shaped structures. Since the priming activity of ATG4B is more efficient than its deconjugation activity, alterations in ATG4B levels were shown to mostly affect deconjugation rather than priming [54]. Our findings using the LC3B biosensor were complementary to this notion, as we were able to observe a stark increase in the number of Aqua-LC3B-II puncta structures when *ATG4B* was silenced. By using a combination of broad and sensitive approaches to quantify FRET, we provided the first proof of concept that proLC3B priming events occur at discrete sites in cells. It is likely that these sites are already present to a lower extend under basal conditions, and they are highlighted only when ATG4B priming activity is altered. With microscopy approaches with higher resolution, it might be possible to reveal the existence of these reservoirs under basal conditions as well. In this light, the LC3B FRET biosensor has the unique capacity to identify these priming reservoirs in living cells and with subcellular resolution, underlining the superior sensitivity of the LC3B biosensor to explore the functional relevance of these structures and the proteins regulating their formation. On the other hand, it should also be noted that the biosensor is specific to ATG4B-LC3B axis, and it should not be intended as a generalized autophagy reporter. Therefore, as with any other reporter, precaution should be taken when interpreting the results reported by the biosensor and understand that the readouts are restricted to ATG4B/LC3B-mediated autophagy.

Furthermore, we confirmed that the isoform ATG4B is the major cysteine protease priming the LC3B biosensor, and that its knockout results in a complete lack of priming. Additionally, we provided evidence that ATG4A could mildly contribute to the priming of the LC3B biosensor in the absence of ATG4B, corroborating previous findings concerning a functional redundancy among these isoforms [24,68]. Interestingly, our biosensor provided novel information on the relevance of specific ATG4B residues for its priming activity. In this light, we observed an unexpected ability of mutant ATG4B^W142A^ to fully prime proLC3B. Trp142 localizes near the catalytic Cys74 residue, and it was suggested to be responsible for LC3 tail recognition [35]. In *in vitro* cleavage assays, ATG4B with mutated Trp142 displayed a significantly reduced ability to cleave C-terminally tagged LC3 [35]. Based on these findings, we were expecting to observe a reduced LC3B priming with ATG4B^W142A^, and no priming was expected with the catalytically-dead mutant ATG4B^C74S^. While ATG4B^C74S^ was incapable of priming proLC3B, we observed a full priming of the LC3B biosensor in the presence of the ATG4B^W142A^ construct. Not only these results corroborate the high efficiency of ATG4B to cleave proLC3B even in conditions where its catalytic activity is severely reduced, but they also highlight a drastic difference between *in vitro* findings and data obtained in more complex paradigms.

Given the rising interest in developing inhibitors that block the early stages of autophagy by targeting ATG4B, we challenged the LC3B biosensor with a selection of available inhibitors. Again, our biosensor demonstrated to be a useful tool to investigate the mode of action and the efficacy of these compounds at the concentrations and timepoints chosen for the analyses. First, we observed increased amounts of Aqua-LC3B-II puncta after the incubation with all the inhibitors, indicating a reduction in the deconjugation activity of ATG4B. These results were not surprising, as the deconjugation activity of ATG4B was reported to be less efficient than the priming, and therefore more prone to get affected upon inhibition [54]. Furthermore, our data also show that none of the inhibitors was able to completely abolish the priming activity of ATG4B towards LC3B. Indeed, we did not observe a cytosolic distribution of the LC3B biosensor, nor ΔLifetime values similar to those measured with the priming-defective G120A biosensor. Despite an incomplete inhibition on priming, we found that the cells treated with NSC or ZPCK exhibited a significant reduction in LC3B priming compared to control cells. Cells treated with MG132 – a proteasome inhibitor with a capacity to inhibit cysteine proteases [55,56] – exhibited a significant increase in the mean ΔLifetime values, along with a positive shift in the histogram mode value. In contrast, they did not display significant amounts of pixels with high ΔLifetime. It is possible that the incubation with MG132 does not prevent the complete degradation of tdLanYFP once this moiety has been cleaved from the biosensor. In this case, the presence of tdLanYFP in the close vicinity of Aqua-LC3B-II puncta would lead to unspecific FRET events, potentially unrelated to the proLC3B priming readout of the biosensor. This is the reason why a multi-parameter FRET quantification – mean ΔLifetime, number of pixels with high ΔLifetime, histogram distribution of the ΔLifetime values – is mandatory to characterize the specificity of ATG4B inhibitors. In this light, we propose that an efficient ATG4B inhibitor should display a significant difference from controls in the three methods of analysis. A compound that did not meet all the criteria but still displayed a significant increase in the number of high-ΔLifetime pixels with a positive histogram mode value shift and showing a band compatible with the unprimed biosensor was FMK-9a. Although FMK-based compounds were shown to be very potent ATG4B inhibitors [30,50,69], a recent study showed that FMK-9a induces autophagy independently of its inhibition on ATG4B activity [51]. Therefore, our findings support these results since FMK-9a did not meet all the criteria to be considered as an efficient ATG4B inhibitor. Finally, our results on Tioconazole and LV-320 indicate that these two compounds inhibit the priming of proLC3B to a lesser extent than the other compounds. Since they only displayed a positive shift in the histogram mode values and did not meet any other criteria, we propose that they should be considered as mild ATG4B inhibitors. Overall, our results underline the lack of inhibitors that can fully inhibit the priming activity of ATG4B. Future screenings using the LC3B biosensor will be useful to identify new inhibitory compounds, as one would expect to observe a FRET behavior similar to that of the G120A biosensor or *ATG4* KO cells in case of a full inhibition of proLC3B priming.

In addition to revealing the differential mode of actions of ATG4 inhibitors, the sensitivity of our biosensor also allowed us to uncover a CDK1-dependent regulation of the ATG4B-LC3B axis at mitosis. Our investigation in mitotic cells indicate that the inhibition of autophagy during cell division is not linked with the accumulation of proLC3B reservoirs. However, the FRET response observed upon CDK1 inhibition in mitotic cells suggests a CDK1-dependent regulation of ATG4B-LC3B nexus. We suggest that this regulation could possibly be regulated by ULK1, one of the mitotic targets of CDK1 [63]. ULK1 is known to be activated at the autophagosome formation site in interphase cells, where it phosphorylates ATG4B to inhibit its catalytic activity towards LC3B [70]. Thus, CDK1 inhibition during mitosis could not only trigger the re-activation of early autophagy targets such as ATG13, ULK1, ATG14 and TFEB [63], but it could also inactivate downstream actors such as ATG4B and LC3B through the direct re-activation of ULK1 which, in turn, inhibits ATG4B.

Since it is possible to calculate the number of pixels with high ΔLifetime, additional information can be provided by localizing these pixels at the subcellular level. We observed a consistent presence of pixels with high ΔLifetime around or on puncta-shaped structures, either upon ATG4B inhibition or *ATG4B* silencing. In this light, we suggest that the local scarcity or the inhibition of ATG4B may cause alterations in the proLC3B priming rates in discrete areas of the autophagosomes, which could be considered as priming “hotspots”. As previously mentioned, these reservoirs or “hotspots” with reduced proLC3B priming rates may be sites where proLC3B is temporarily stored while trying to re-establish the full priming capacity of ATG4B.

Overall, we present the LC3B biosensor as a second-generation FRET biosensor that can report on the regulation of the soluble and the lipidated forms of LC3B by ATG4B. First, this tool can be used to infer on the structural properties of ATG4B and on its enzymatic activity. Thanks to its dual FRET/localization readout, it can also be used to follow LC3B priming and turnover with superior spatiotemporal resolution. Finally, the LC3B biosensor has the potential to be used in high-content screenings to identify more potent ATG4B inhibitors and reveal their mode of action in living cells, which is a unique feature of the biosensor compared to *in vitro* screening methodologies. Thus, the LC3B biosensor paves the way to ATG4B-targeted therapies in complex diseases.

## Materials and Methods

### Expression vectors and molecular cloning

All the plasmids used in this study are listed in Supplementary Table 1. The cloning reactions were performed using the Gibson Assembly Master Mix (New England Biolabs). Site-directed mutagenesis was performed with the Quik-Change kit (Agilent). All the constructs from cloning and mutagenesis reactions were verified using a 3130 XL sequencer (Applied Biosystems) and a BigDye Terminator V3.1 sequencing kit (Applied Biosystems).

### Cell culture and transfections

U2OS cells (HTB-96) were purchased from American Type Culture Collection. Control and *ATG4* KO HeLa cells were kind gifts of Dr. Robin Ketteler (UCL, LMCB, United Kingdom). Cells were cultured in DMEM (Thermo Fisher Scientific) supplemented with 10% FBS (Eurobio Scientific) and penicillin-streptomycin (100 U/mL, Thermo Fisher Scientific) and maintained at 37°C with 5% CO2. All cell lines were routinely checked for the absence of mycoplasma. Before imaging, normal growth media was replaced with phenol red-free Leibovitz’s L-15 medium (Thermo Fisher Scientific) supplemented with 20% FBS and penicillin-streptomycin (100 U/mL). Cells were seeded at 70% confluence in Nunc Lab-Tek II Chamber slides (Thermo Fisher Scientific) or Cellview cell culture slides (Greiner bio-one, 543979) for live cell imaging, 24-well plates for immunocytochemistry, or 6-well plates for total cell lysates. Plasmid DNA transfection, or plasmid DNA and siRNA co-transfection experiments were performed using Lipofectamine 2000 (Invitrogen) according to the manufacturer’s instructions. Cells were analyzed 48h after transfection. AllStars negative control siRNA (SI03650318) and the *ATG4B*-specific siRNA (SI03156314) were purchased from QIAGEN.

### Chemical compounds

The chemical compounds used in this study were as follows: Bafilomycin A1 (Sigma-Aldrich, B1793), FMK 9a (MedChemExpress, HY-100522), LV-320 (MedChemExpress, HY-112711), MG-132 (Selleckchem, S2619), Nocodazole (Sigma-Aldrich, M1404), NSC 185058 (Selleckchem, S6716), RO-3306 (Sigma-Aldrich, SML0569), Tioconazole (Sigma-Aldrich, 03907), Torin1 (Sigma-Aldrich, 475991), *Z-L-Phe* chloromethyl ketone (Sigma-Aldrich, 860794). All chemical compounds were dissolved in dimethyl sulfoxide (Sigma-Aldrich, D2438) and stored at −80°C. For starvation assay, a home-made Hank’s Balanced Salt Solution (HBSS) containing 8 mg/ml NaCl, 0.4 mg/ml KCl, 0.06 mg/ml KH_2_PO_4_, 0.048 mg/ml Na_2_HPO_4_ anhydrous, 1 mg/ml glucose, 0.348 mg/ml NaHCO_3_ and penicillin-streptomycin (100 U/mL) was used. Concentrations and durations of each treatment are indicated in the figure legends.

### Western blotting

To collect total cell lysates, cells were rinsed with ice-cold Phosphate Buffer Saline (PBS) (Euromedex, ET330-A) and lysed on ice in a buffer containing 50 mM Tris-HCL (pH 7.4), 150 mM NaCl, 1% Triton X-100 (Euromedex, 2000-A), 1.5 mM MgCl_2_, supplemented with 0.2 mM Na_3_VO_4_, 0.5 mM DTT (Thermo Fisher Scientific, R0861), 4 mg/ml NaF, 5.4 mg/ml β-glycerolphosphate and a protease inhibitor cocktail (Roche, 11873580001) immediately prior to lysis. Lysates were centrifuged at 13000 *g* for 20 minutes at 4°C. Protein levels were quantified by using the Bradford protein assay dye reagent (BioRad, 5000006). Lysates were then heated in Laemmli sample buffer at 95°C for 5 minutes, resolved in home-made Acrylamide/Bis 37.5:1 SDS-PAGE mini gels and transferred onto nitrocellulose membrane (Amersham™ Protran®, 10600004). Membranes were blocked in a solution containing 5% skimmed milk in TBS-T (TBS [Euromedex, ET220] containing 0.1% Tween [Euromedex, 2001-B]) and incubated overnight at 4°C with primary antibodies diluted in the blocking solution. The next day, membrane was washed in TBS-T, incubated with the secondary antibody diluted in the blocking solution for 1h at room temperature, and washed again in TBS-T prior to detection. The primary antibodies and dilutions were as follows: rabbit anti-Actin (Sigma-Aldrich, A5060; 1:1000), ATG4B (Cell Signaling, 5299; 1:1000), LC3B (Cell Signaling, 3868; 1:1000). The secondary antibody used was a horseradish-peroxidase-conjugated goat anti-rabbit antibody (Jackson ImmunoResearch; 1:6000-1:10000). After incubating the membrane in an ECL western blotting substrate (Thermo Fisher Scientific, 32209), chemiluminescence signals were captured on a film (Thermo Fisher Scientific, 34091) and developed with a CURIX 60 developer (Agfa Healthcare). The density of the bands was quantified by using the *Gel Analyzer* function in Fiji (NIH) software. The relative abundance of each band was calculated by normalizing the density of the band to that of the respective loading control.

### Immunocytochemistry, confocal and FLIM microscopy

For immunocytochemistry, cells were seeded on 15 mm round coverslips placed onto 24-well plates. Cells were washed with 1X PBS and fixed in 1X PBS containing a mixture of 4% paraformaldehyde (Electron Microscopy Sciences, 15710) and 0.2% Glutaraldehyde (Euromedex, EM-16221) at room temperature for 20 minutes. After washing in 1X PBS, cells were permeabilized with 0.2% Triton in PBS for 10 minutes, washed again in 1X PBS and blocked for 1h in 5% BSA (Euromedex, 04-100-812-C) in 1X PBS at room temperature. Cells were incubated overnight at 4°C with primary antibodies diluted in the blocking buffer, and then washed with 1X PBS. Cells were then incubated with the secondary antibody diluted in the blocking buffer for 45 minutes at room temperature. Primary monoclonal anti-LAMP2 (Abcam, ab25631; 1:200) was used as a primary antibody and a goat anti-mouse IgG (H+L) cross-adsorbed antibody Alexa Fluor™ 647 (Thermo Fisher Scientific, A-21235; 1:500) was used as a secondary antibody. After washing in 1X PBS, coverslips were mounted in ProLong Gold Antifade reagent (Invitrogen, P36930). Cells were imaged with a Leica SP8 inverted confocal microscope equipped with a 63x oil immersion objective (NA 1.4). Aquamarine fluorescence was acquired with a 440 nm excitation laser, and an emission wavelength of 467-499 nm. The fluorescence of tdLanYFP and of LAMP2/Alexa 647 were captured by using a 514 nm and a 633 nm argon laser, respectively. The emission wavelengths were 525-565 nm for tdLanYFP, and 650-720 nm for LAMP2/Alexa 647. For FLIM analyses, images were acquired with a time-gated custom-made setup based on a spinning disk microscope as described in [71]. Aquamarine was used as a FRET donor in all experiments, and excited at 440 ± 10 nm with a supercontinuum picosecond pulsed laser source. Emission was selected using a band pass filter of 483/35 nm. The FLIM setup was controlled by the Inscoper Suite solution (Inscoper, France), and Aquamarine lifetime was measured in real-time during acquisition with the Inscoper software.

### Image analysis

All the image analysis were performed in Fiji software. 3D puncta counting and fluorescence colocalization analyses illustrated in Fig. 1B-D and Fig. S1 were performed by using the macro developed by Cordelières and Zhang [72] in batch processing mode, and available in a GitHub repository at https://github.com/NEUBIAS/neubias-springer-book-2020. The minimum size of the objects for Aquamarine-LC3B and LAMP2/Alexa 647 was set to 10 voxels. The threshold to separate the objects from the background was set manually for both channels. The total number of objects in Aquamarine-LC3B channel was used to determine the number of Aqua-LC3B-II puncta-shaped structures. The objects in the Aquamarine-LC3B channel superposing with the LAMP2/Alexa 647 objects were used for colocalization analyses, and only the Aquamarine-LC3B objects superposing with the LAMP2/Alexa 647 objects with a ratio of 0.5 or more were quantified for analyses. The colocalizing objects were then normalized to the total number of Aquamarine-LC3B objects. For FLIM analysis, mean ΔLifetime values were calculated as previously described [38]. In all experiments, Aquamarine lifetime was calculated by the Inscoper software only when the pixel-by-pixel fluorescence intensity in the first gate was above 1000 grey levels. The number of Aqua-LC3B-II puncta structures in the accompanying fluorescence images (Fig. 2A, S2A, 5A-F, S9A-B) were quantified using the *Find Maxima* function in the Fiji imaging software, and by setting the prominence value as 1500. To analyze the high-ΔLifetime pixels, the *Histogram* tool in Fiji was used to measure the number of pixels with a lifetime between 2000 and 4000 psec. Each histogram was then converted to a ΔLifetime format by using the mean lifetime value of the donor-only construct as a normalizer. To determine the number of pixels with high ΔLifetime, the mean ΔLifetime value of the G120A biosensor or the mean ΔLifetime value of the WT biosensor expressed in *ATG4B* SKO cells were used as a threshold. The number of pixels showing G120A biosensor-like ΔLifetime or higher were then quantified and normalized to the total number of pixels, and this to determine the high-ΔLifetime pixel ratio per cell. For line analysis, a 17.8 μm linear region of interest (ROI) that contains both the high- and low-ΔLifetime pixels was manually drawn near or on the puncta-like structures. The *Plot profile* function in Fiji was then used to obtain ΔLifetime values on the drawn line, which were then plotted. For histogram analyses, the average number of pixels per ΔLifetime was quantified for each condition.

### Statistical analysis

All statistical tests were performed by using GraphPad Prism 9. Two-way ANOVA with Tukey method was applied to make multiple comparisons in the following figures: 1C-D; 2B-C, F; 3B-C; 4B-C; 5B-F; 6B-D; 7B-C; S2B, F; S3B-C; S5A-B: S8B-C: S9B; S10B. Two-way ANOVA with two-stage step-up method of Benjamini, Krieger and Yekutieli was applied to make multiple comparisons in the following figures: 1F-G, S2D and S5D. Correlation analysis between the ΔLifetime values and the puncta numbers were performed to compute R^2^ and P values in Fig. S6.

### Figure preparation

The cartoon in Figure 1A was prepared by using the illustrations available at https://smart.servier.com/ [73]. Graphs and figures were assembled in GraphPad Prism 9 and Inkscape, respectively.

## Abbreviations

ATG: autophagy-related
AURKA: Aurora kinase A
BafA_1_: Bafilomycin A_1_
CDK1: Cyclin-dependent kinase 1
DKO: double knockout
FLIM: fluorescence lifetime imaging microscopy
FRET: Förster’s resonance energy transfer
GABARAP: gamma-aminobutyric acid (GABA) type A receptor-associated protein
HBSS: Hank’s balanced salt solution
KO: knockout
LAMP2: lysosomal associated membrane protein 2
MAP1LC3/LC3: microtubule-associated protein 1 light chain 3
mTORC1: mammalian target of rapamycin complex 1
NSC: NSC 185058
PE: phosphatidylethanolamine
SKO: single knockout
TFEB: Transcription Factor EB
TKO: triple knockout
ULK1: Unc-51 like autophagy activating kinase 1
ZPCK: *Z-L*-phe chloromethyl ketone

## Data and material availability

Plasmids and macro used in this study and the source data that support the findings are available from the corresponding authors (G.B. and M.T.) on request.

## Acknowledgments

We thank P. Govindin (MetaGenoPolis, INRAe, Jouy-en-Josas, France) for preliminary experiments with the LC3B biosensor, S. Dutertre and X. Pinson at the Microscopy Rennes Imaging Center (MRic, *Biologie, Santé, Innovation Technologique* - BIOSIT, Rennes, France) and G. Le Marchand (IGDR, Rennes, France) for help and assistance. MRic is member of the national infrastructure France-BioImaging supported by the French National Research Agency (ANR-10-INBS-04). We also thank R. Ketteler (UCL, LMCB, United Kingdom) for sharing pGEX GST-ATG4B plasmid and *ATG4* KO HeLa cells. We are grateful to S. Ley-Ngardigal, R. Smith, C. Chapuis and S. Zentout for technical assistance with the experiments, and Ç. Tuna for help with the image analysis. This work was supported by the *Centre National de la Recherche Scientifique* (CNRS), the University of Rennes 1, the *Ligue Contre le Cancer Comité d’īlle et Vilaine et du Finistère* and the *Association pour la Recherche sur le Cancer* (ARC) to G.B., and by the *Institut National du Cancer* (INCa) and *ITMO Cancer/Aviesan* to M.T. E.B.G. was supported by a fellowship from the *Ligue Contre le Cancer* and *Région Bretagne* (Brittany region, France).

## Author Contributions

E.B.G. designed, performed and analyzed the experiments and wrote the manuscript; A.C. performed the experiments and revised the manuscript, M.T. co-supervised the work, revised the manuscript and provided funding; G.B. co-supervised the work, designed the experiments, edited and revised the manuscript, and provided funding.

## Conflict of interest

The authors declare no conflict of interest.

## Notes

### Competing Interest Statement

The authors have declared no competing interest.

### Summary of Updates

Addition of Fig. 7 and of several supplemental data

