## Supplementary Information v2 for "The LC3B FRET biosensor monitors the modes of action of ATG4B during autophagy in living cells"

**Elif Begüm Gökerküçük et al.**

#### Supplementary figures

**Figure S1.** The G120A LC3B biosensor is not sensitive to autophagy induction and/or lysosomal inhibition, and it does not colocalize with LAMP2. Representative fluorescence images of U2OS expressing the G120A biosensor and stained for endogenous LAMP2. To investigate the changes in Aqua-LC3B puncta numbers and their colocalization with LAMP2, cells were treated with: DMSO (6h), BafA1 (6h, 100 nM), Torin1 (3h, 250 nM), Torin1 (3h, 250 nM) + BafA1 (6h, 100 nM), HBSS (1h), HBSS (1h) + BafA1 (6h, 100 nM). Scale bar: 9 µm.

**Figure S2.** The WT or G120A donors do not respond to *ATG4B* silencing. (**A**) Representative fluorescence and ΔLifetime images of U2OS cells co-expressing the WT or G120A donor with control or *ATG4B*-specific siRNAs, and analyzed by FRET/FLIM. Squares on the top images of WT or G120A donor panels illustrate the location of the enlarged images. Dotted lines on the enlarged images illustrate where the line analysis was performed. Pseudocolor scale: pixel-by-pixel ΔLifetime. Scale bars: overviews, 40 µm; enlarged, 6 µm. (**B**) Mean ΔLifetime analysis of U2OS cells co-expressing the WT or G120A donor with control or *ATG4B* siRNA. Representative western blotting images (**C**) and corresponding quantifications (**D**) of total lysates from U2OS cells co-expressing the WT or G120A biosensor with control or *ATG4B*-specific siRNAs. IB1 and IB2 correspond to the same lysates blotted for overexpressed (IB1) or endogenous (IB2) LC3B forms. Loading control: Actin. $n$ = 3 independent experiments. (**E**) Line and (**F**) number of Aqua-LC3B-II puncta analyses of U2OS cells co-expressing the WT or G120A donor with control or *ATG4B* siRNA. $n$ = 10 cells per condition from one representative experiment (of three) in (**B**) and (**F**). *$P < 0.05$, ****$P < 0.0001$, ns (not significant) as determined by two-way ANOVA with Tukey's multiple comparison test in (**B**) and (**F**), and with two-stage step-up method of Benjamini, Krieger and Yekutieli's multiple comparison test to control the false discovery rate in (**D**).

**Figure S3.** No FRET is detected when donor-bound LC3B and acceptor-bound LC3B are co-expressed. (**A**) Representative fluorescence and ∆Lifetime images of U2OS cells co-expressing the WT or G120A biosensor, or donor (Aquamarine-proLC3B) + acceptor (proLC3B-tdLanYFP) constructs with control or *ATG4B*-specific siRNAs, and analyzed by FRET/FLIM. Squares on the top images of WT or G120A biosensor, or donor + acceptor panels illustrate the location of the enlarged images. Dotted lines on the enlarged images illustrate where the line analysis was performed. Pseudocolor scale: pixel-by-pixel ∆Lifetime. Scale bars: overviews, 40 µm; enlarged, 6 µm. Mean ∆Lifetime (**B**), number of high ∆Lifetime pixels (**C**), line (**D**), and histogram (**E**) analyses of U2OS cells co-expressing the WT or G120A biosensor, or donor + acceptor constructs with control or *ATG4B*-specific siRNAs in (**B**), (**D**), and (**E**), and the WT donor, biosensor or donor + acceptor constructs with control or *ATG4B* siRNA in (**C**). Vertical dotted lines on each histogram depict the mode value in (**E**). $n = 10$ cells per condition from one representative experiment (of three) in (**B**), (**C**), and (**E**). *$P < 0.05$, **$P < 0.01$, ****$P < 0.0001$, ns (not significant) as determined by two-way ANOVA with Tukey's multiple comparison test in (**B**) and (**C**).

**Figure S4.** The LC3B biosensor and endogenous LC3B proteins cannot be primed in cells lacking ATG4B. Representative western blotting images of total lysates from HeLa control, *ATG4B* SKO, *ATG4A/B* DKO, *ATG4A/B/C* TKO cells expressing the WT or G120A biosensor. IB1 and IB2 correspond to the same lysates blotted for overexpressed (IB1) or endogenous (IB2) LC3B forms. Loading control: Actin. *n* = 3 independent experiments.

**Figure S5.** The ectopic expression of active ATG4B does not alter LC3B donor-only ∆lifetime variations in *ATG4B* SKO cells. Mean ∆Lifetime (**A**) and number of high ∆Lifetime pixels (**B**) analyses of control and *ATG4B* SKO HeLa cells co-expressing the WT donor with an empty vector, or with vectors expressing WT ATG4B, ATG4B$^{C74S}$ or ATG4B$^{W142A}$. (**C**) Histogram analysis of HeLa control cells co-expressing the WT biosensor with an empty vector, or with vectors expressing WT ATG4B, ATG4B$^{C74S}$ or ATG4B$^{W142A}$. Vertical dotted lines on each histogram depict the mode value. *n* = 10 cells per condition from one representative experiment (of three) in (**A**), (**B**) and (**C**). (**D**) Representative western blotting images and corresponding ATG4B/Actin quantification of total lysates from HeLa control and *ATG4B* SKO cells co-expressing the WT biosensor with an empty vector, or with vectors expressing WT ATG4B, ATG4B$^{C74S}$ or ATG4B$^{W142A}$. IB1 and IB2 correspond to the same lysates blotted for overexpressed (IB1) or endogenous (IB2) LC3B forms. Short and long ATG4B exposures are shown. Loading control: Actin. *n* = 3 independent experiments. ns (not significant) as determined by two-way ANOVA with Tukey's multiple comparison test in (**A**) and (**B**), and with two-stage step-up method of Benjamini, Krieger and Yekutieli's multiple comparison test to control the false discovery rate in (**D**).

**Figure S6.** The number of Aqua-LC3B-II puncta and the mean ∆Lifetime values do not correlate. Correlation analyses of Aqua-LC3B-**II** puncta and the mean ∆Lifetime values of HeLa cells expressing the WT biosensor and treated with DMSO (6h), MG132 (6h, 1 µM), Tioconazole (6h, 4 µM), LV-320 (6h, 120 µM), FMK-9a (6h, 10 µM), NSC 185058 (6h, 100 µM), or *Z*-*L*-Phe chloromethyl ketone (6h, 3 µM). $n$ = 10 cells per condition from one representative experiment (of three).

**Figure S7.** The treatment with FMK-9a, NSC and ZPCK results in the accumulation of unprimed biosensor. Representative western blotting and corresponding Aqua-proLC3B-tdLanYFP/Actin quantification of total lysates of HeLa cells expressing the WT or G120A biosensor and treated with DMSO (6h), or expressing the WT biosensor and treated with Tioconazole (6h, 4 µM), LV-320 (6h, 120 µM), FMK-9a (6h, 10 µM), NSC 185058 (6h, 100 µM), or *Z*-*L*-Phe chloromethyl ketone (6h, 3 µM). IB1 and IB2 correspond to the same lysates blotted for overexpressed (IB1) or endogenous (IB2) LC3B forms. Aqua-proLC3B-tdLanYFP/Actin quantification is performed by normalizing the density of Aqua-proLC3B-tdLanYFP band in IB1 to that of the respective loading control in IB2. Loading control: Actin. *n* = 3 independent experiments.

**Figure S8.** FRET/FLIM data analyzed by using sensitive methods reveal the mode of action of Tioconazole and LV-320 in cells. (**A**) Representative fluorescence and ∆Lifetime images of HeLa cells expressing the WT or G120A biosensor, treated with DMSO (6h), and analyzed by FRET/FLIM. Representative fluorescence and ∆Lifetime images of HeLa cells expressing the WT biosensor and treated with the following compounds: Tioconazole (6h, 4 µM) (**B**), LV-320 (6h, 120 µM) (**C**). Squares on the top images of WT or G120A biosensor panels illustrate the location of the enlarged images. Dotted lines on the enlarged images illustrate where the line analysis was performed. Pseudocolor scale: pixel-by-pixel ∆Lifetime. Scale bars: overviews, 40 µm; enlarged, 6 µm. Number of high ∆Lifetime pixels analysis of HeLa cells expressing the WT donor or biosensor and treated with Tioconazole (6h, 4 µM) (**B**) or LV-320 (6h, 120 µM) (**C**). Line, and histogram analyses of HeLa cells expressing the WT or G120A biosensor and treated with Tioconazole (6h, 4 µM) (**B**) or LV-320 (6h, 120 µM) (**C**). *n* = 10 cells per condition from one representative experiment (of three) in (**B, C**). ns (not significant) as determined by two-way ANOVA with Tukey's multiple comparison test in (**B, C**).

**Figure S9.** The LC3B biosensor reveals the mode of action of MG132 in cells. (**A and B**) Representative fluorescence and ∆Lifetime images of HeLa cells expressing the WT or G120A biosensor, treated with DMSO (6h) or MG132 (6h, 1 µM), and analyzed by FRET/FLIM. Pseudocolor scale: pixel-by-pixel ∆Lifetime. Scale bars: overviews, 40 µm; enlarged, 6 µm. Mean ∆Lifetime (**B**), number of Aqua-LC3B-II puncta (**B**), histogram (**C**), and line (**D**) analyses of HeLa cells expressing the WT or G120A biosensor and treated with DMSO (6h) or MG132 (6h, 1 µM). Number of high ∆Lifetime pixels analysis (**B**) of HeLa cells expressing the WT donor or biosensor and treated with DMSO (6h) or MG132 (6h, 1 µM). Vertical dotted lines on each histogram depict the mode value in (**C**). *n* = 10 cells per condition from one representative experiment (of three) in (**B**) and (**C**). (**E**) Representative western blotting of total lysates of HeLa cells expressing the WT or G120A biosensor and treated with DMSO (6h), or expressing the WT biosensor and treated with MG132 (6h, 1 µM). IB1 and IB2 correspond to the same lysates blotted for overexpressed (IB1) or endogenous (IB2) LC3B forms. Loading control: Actin. *n* = 3 independent experiments. ***P* < 0.01, *****P* < 0.0001, ns (not significant) as determined by two-way ANOVA with Tukey's multiple comparison test in (**B**).

**Figure S10.** The FRET readout of the AURKA biosensor revealed a lowered activation of AURKA in cells treated with RO-3066**.** (**A**) Representative fluorescence and ΔLifetime images of U2OS cells expressing the AURKA biosensor (tdLanYFP-AURKA-Aquamarine) and treated with nocodazole-only (16h, 100 ng/ml) or co-treated with nocodazole (16h, 100 ng/ml) and RO-3306 (2h, 2 µM), and analyzed by FRET/FLIM. Pseudocolor scale: pixel-by-pixel ΔLifetime. Scale bars: 20 µm. (**B**) Mean ΔLifetime analysis of U2OS cells expressing the AURKA donor-only (AURKA-Aquamarine) or AURKA biosensor (tdLanYFP-AURKA-Aquamarine), and co-treated with nocodazole (16h, 100 ng/ml) and DMSO, or with nocodazole (16h, 100 ng/ml) and RO-3306 (2h, 2 µM). $n = 10$ cells per condition from one representative experiment (of three) in (**B**). *$P < 0.05$, ****$P < 0.0001$, ns (not significant) as determined by two-way ANOVA with Tukey's multiple comparison test in (**B**).

### Figure S1

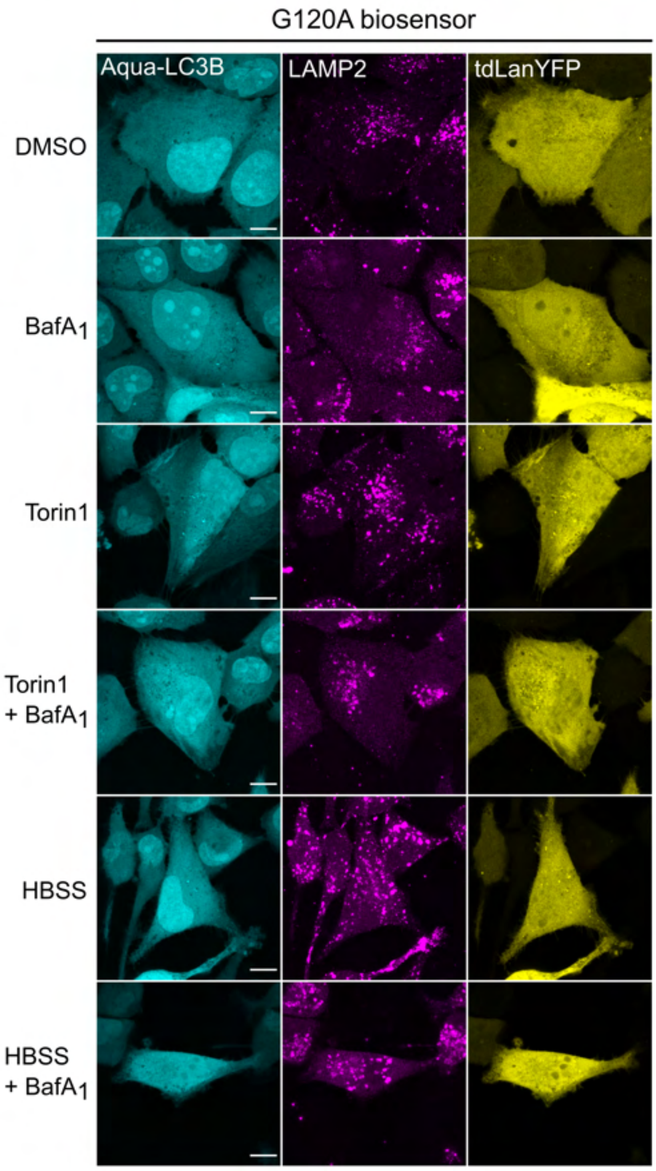
G120A biosensor
Aqua-LC3B
LAMP2
tdLanYFP
DMSO
BafA1
Torin1
Torin1
+ BafA1
HBSS
HBSS
+ BafA1

### Figure S2

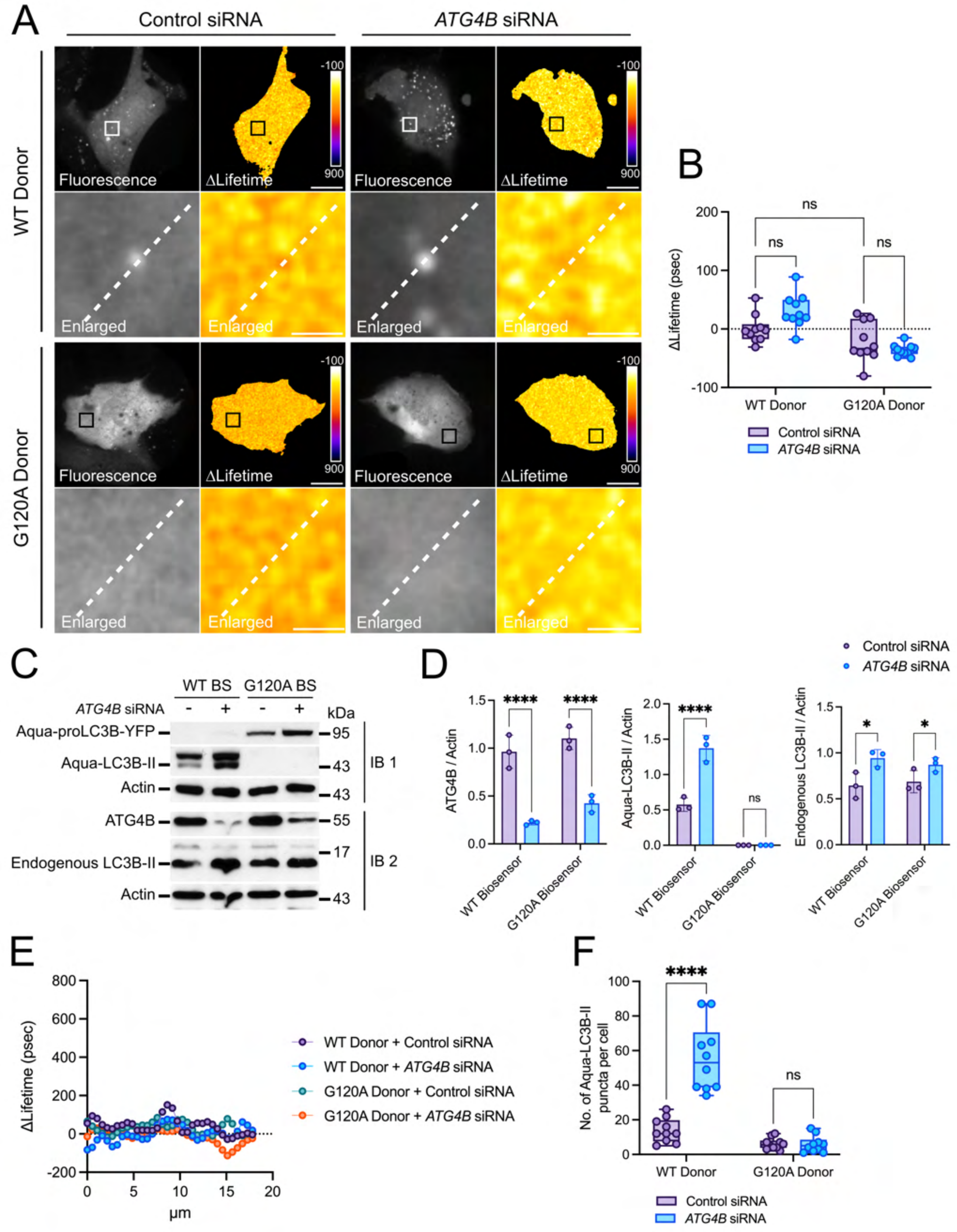

A
Control siRNA
ATG4B siRNA
WT Donor
G120A Donor
Fluorescence
ΔLifetime
Enlarged
-100
900
B
ΔLifetime (psec)
ns
WT Donor
G120A Donor
Control siRNA
ATG4B siRNA
C
WT BS
G120A BS
ATG4B siRNA
kDa
Aqua-proLC3B-YFP
Aqua-LC3B-II
Actin
ATG4B
Endogenous LC3B-II
IB 1
IB 2
95
43
55
17
D
ATG4B / Actin
Aqua-LC3B-II / Actin
Endogenous LC3B-II / Actin
WT Biosensor
G120A Biosensor
****
*
E
ΔLifetime (psec)
μm
WT Donor + Control siRNA
WT Donor + ATG4B siRNA
G120A Donor + Control siRNA
G120A Donor + ATG4B siRNA
F
No. of Aqua-LC3B-II puncta per cell
WT Donor
G120A Donor
Control siRNA
ATG4B siRNA

### Figure S3

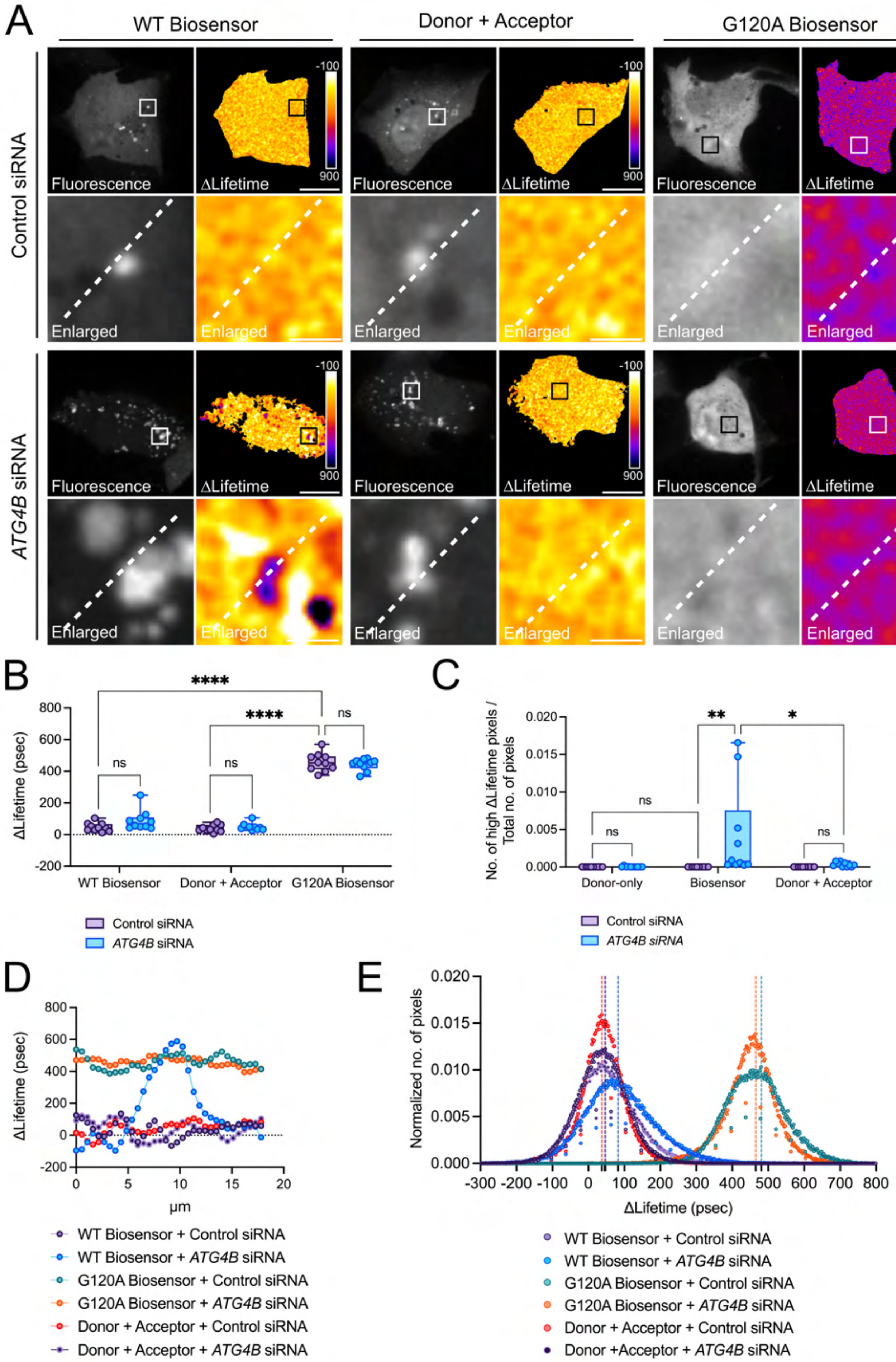
A
WT Biosensor
Donor + Acceptor
G120A Biosensor
Control siRNA
ATG4B siRNA
Fluorescence
ΔLifetime
Enlarged
-100
900
B
****
****
ns
ns
ns
ΔLifetime (psec)
800
600
400
200
0
-200
WT Biosensor
Donor + Acceptor
G120A Biosensor
Control siRNA
ATG4B siRNA
C
No. of high ΔLifetime pixels / Total no. of pixels
0.020
0.015
0.010
0.005
0.000
**
*
ns
ns
ns
Donor-only
Biosensor
Donor + Acceptor
Control siRNA
ATG4B siRNA
D
ΔLifetime (psec)
µm
0
5
10
15
20
WT Biosensor + Control siRNA
WT Biosensor + ATG4B siRNA
G120A Biosensor + Control siRNA
G120A Biosensor + ATG4B siRNA
Donor + Acceptor + Control siRNA
Donor + Acceptor + ATG4B siRNA
E
Normalized no. of pixels
0.020
0.015
0.010
0.005
0.000
-300 -200 -100 0 100 200 300 400 500 600 700 800
ΔLifetime (psec)
WT Biosensor + Control siRNA
WT Biosensor + ATG4B siRNA
G120A Biosensor + Control siRNA
G120A Biosensor + ATG4B siRNA
Donor + Acceptor + Control siRNA
Donor +Acceptor + ATG4B siRNA

### Figure S4

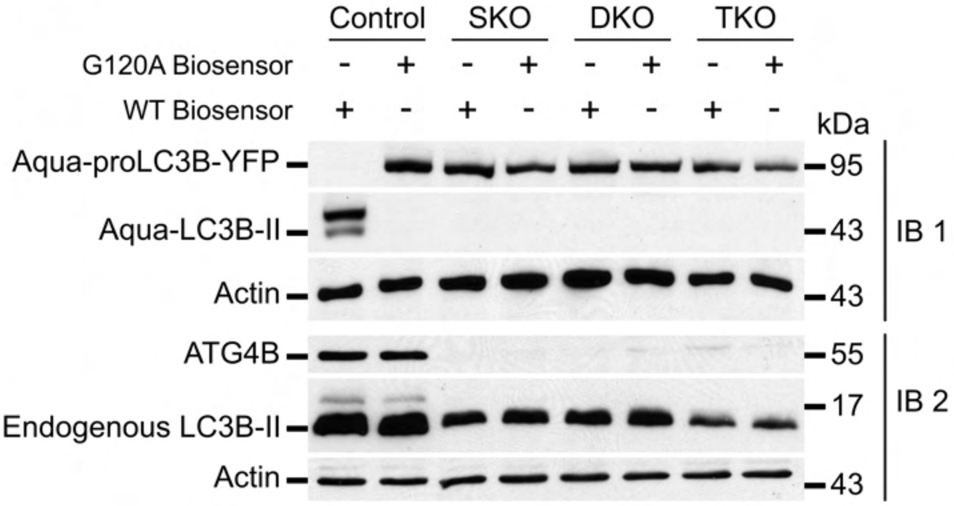
Control
SKO
DKO
TKO
G120A Biosensor
WT Biosensor
kDa
Aqua-proLC3B-YFP
Aqua-LC3B-II
Actin
ATG4B
Endogenous LC3B-II
Actin
95
43
43
55
17
43
IB 1
IB 2

### Figure S5

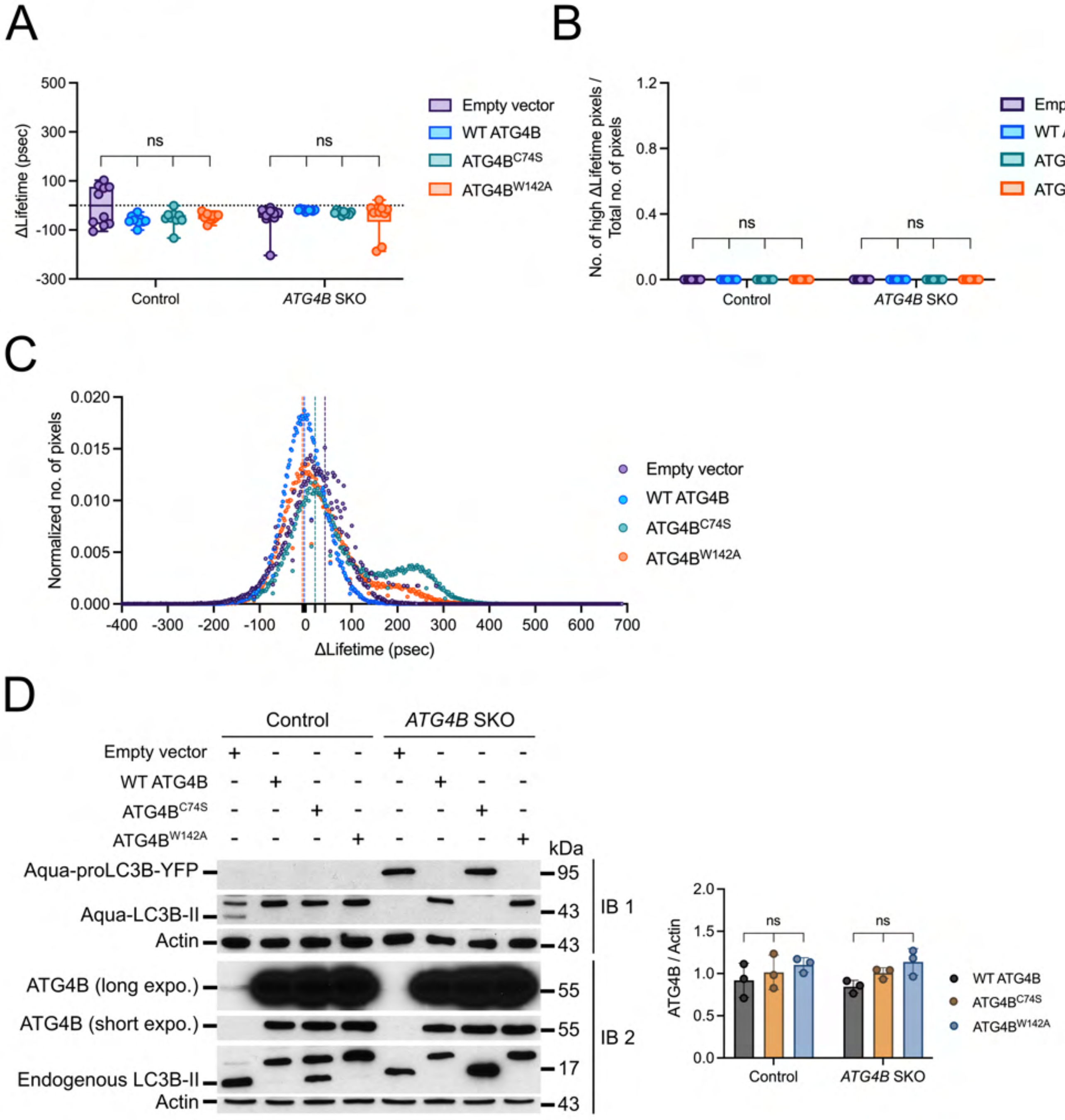
A
ΔLifetime (psec)
500
300
100
-100
-300
ns
ns
Control
ATG4B SKO
Empty vector
WT ATG4B
ATG4B C74S
ATG4B W142A
B
No. of high ΔLifetime pixels / Total no. of pixels
1.2
0.8
0.4
0.0
ns
ns
Control
ATG4B SKO
Empty vector
WT ATG4B
ATG4B C74S
ATG4B W142A
C
Normalized no. of pixels
0.020
0.015
0.010
0.005
0.000
-400
-300
-200
-100
0
100
200
300
400
500
600
700
ΔLifetime (psec)
Empty vector
WT ATG4B
ATG4B C74S
ATG4B W142A
D
Control
ATG4B SKO
Empty vector
WT ATG4B
ATG4B C74S
ATG4B W142A
kDa
Aqua-proLC3B-YFP
Aqua-LC3B-II
Actin
ATG4B (long expo.)
ATG4B (short expo.)
Endogenous LC3B-II
Actin
95
43
43
55
55
17
43
IB 1
IB 2
ATG4B / Actin
2.0
1.5
1.0
0.5
0.0
ns
ns
Control
ATG4B SKO
WT ATG4B
ATG4B C74S
ATG4B W142A

### Figure S6

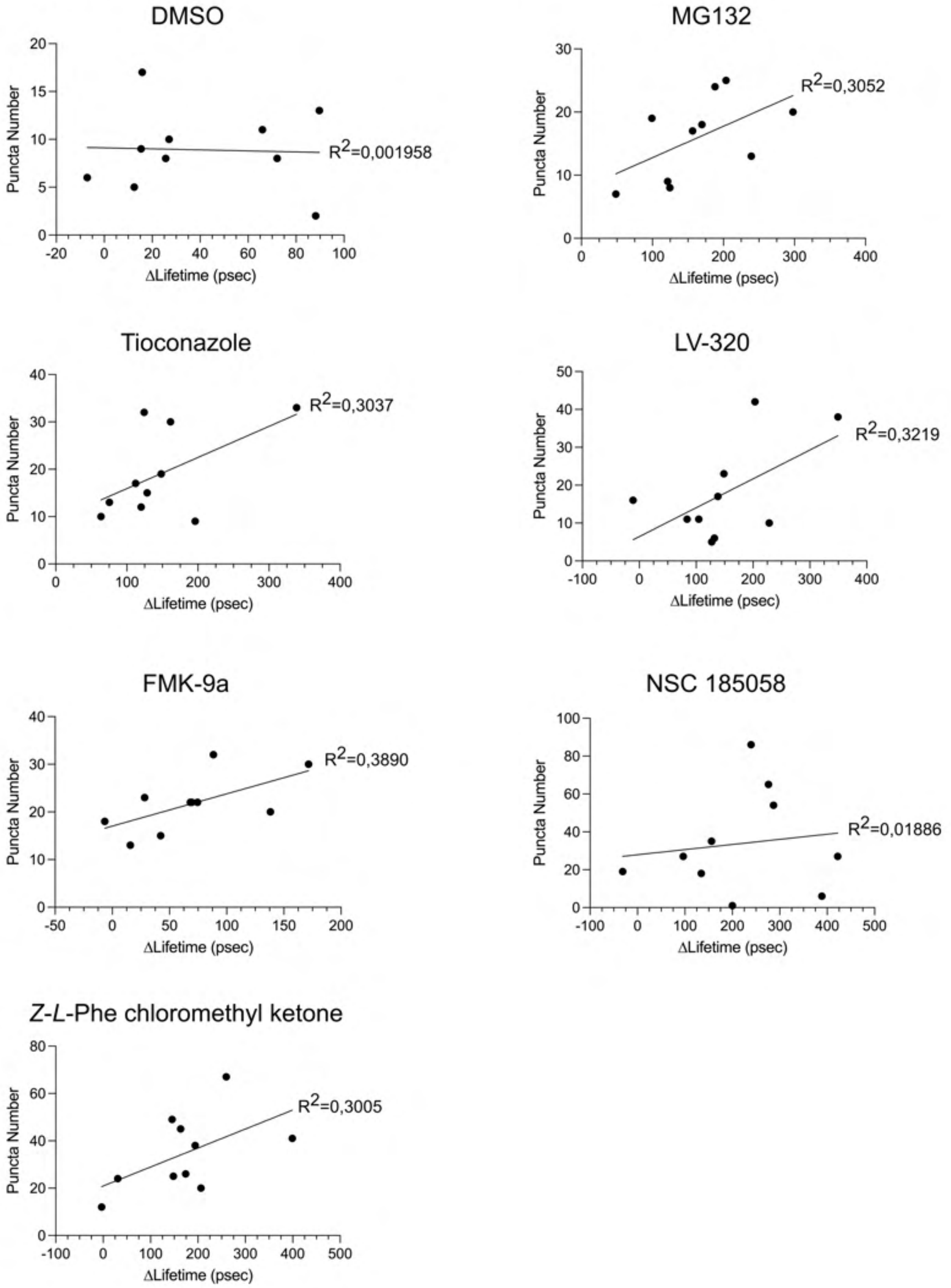

DMSO
Puncta Number
ΔLifetime (psec)
R2=0,001958
MG132
Puncta Number
ΔLifetime (psec)
R2=0,3052
Tioconazole
Puncta Number
ΔLifetime (psec)
R2=0,3037
LV-320
Puncta Number
ΔLifetime (psec)
R2=0,3219
FMK-9a
Puncta Number
ΔLifetime (psec)
R2=0,3890
NSC 185058
Puncta Number
ΔLifetime (psec)
R2=0,01886
Z-L-Phe chloromethyl ketone
Puncta Number
ΔLifetime (psec)
R2=0,3005

### Figure S7

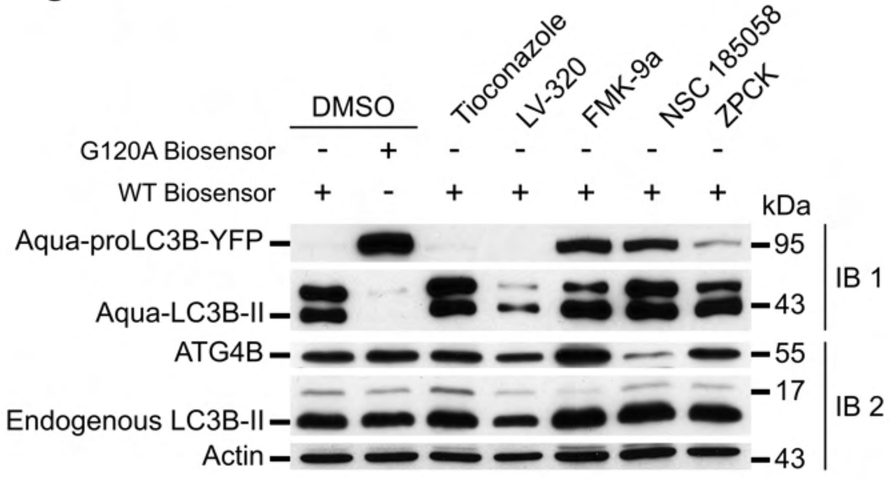

DMSO
Tioconazole
LV-320
FMK-9a
NSC 185058
ZPCK
G120A Biosensor
WT Biosensor
kDa
Aqua-proLC3B-YFP
Aqua-LC3B-II
ATG4B
Endogenous LC3B-II
Actin
95
43
55
17
43
IB 1
IB 2


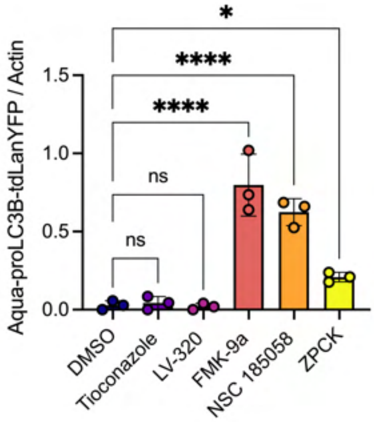

Aqua-proLC3B-tdLanYFP / Actin
1.5
1.0
0.5
0.0
*
****
****
ns
ns
DMSO
Tioconazole
LV-320
FMK-9a
NSC 185058
ZPCK

### Figure S8

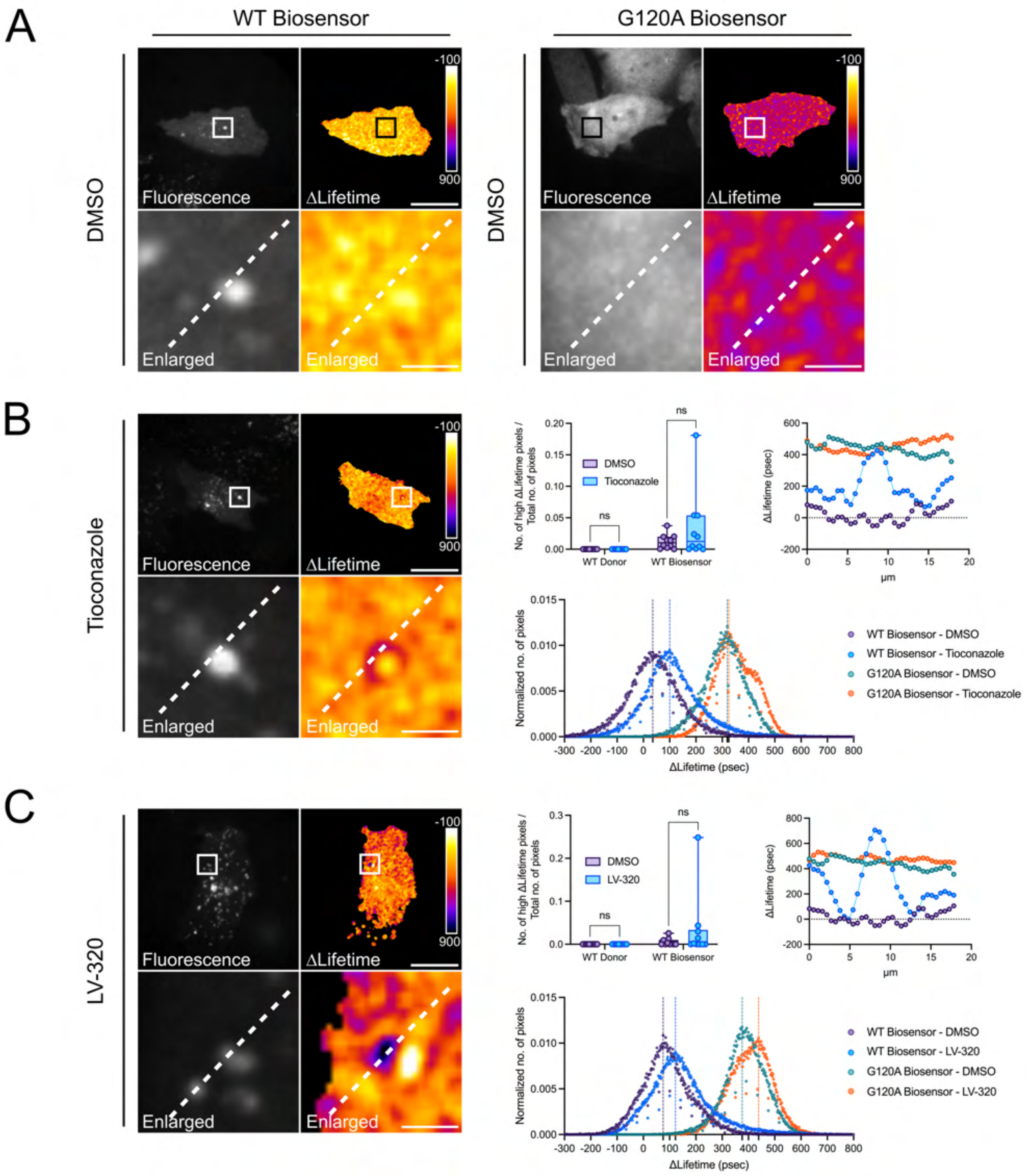

A
WT Biosensor
G120A Biosensor
DMSO
Fluorescence
ΔLifetime
Enlarged
-100
900
B
Tioconazole
ns
DMSO
Tioconazole
WT Donor
WT Biosensor
No. of high ΔLifetime pixels / Total no. of pixels
ΔLifetime (psec)
μm
Normalized no. of pixels
WT Biosensor - DMSO
WT Biosensor - Tioconazole
G120A Biosensor - DMSO
G120A Biosensor - Tioconazole
C
LV-320
WT Biosensor - LV-320
G120A Biosensor - LV-320

Figure S9

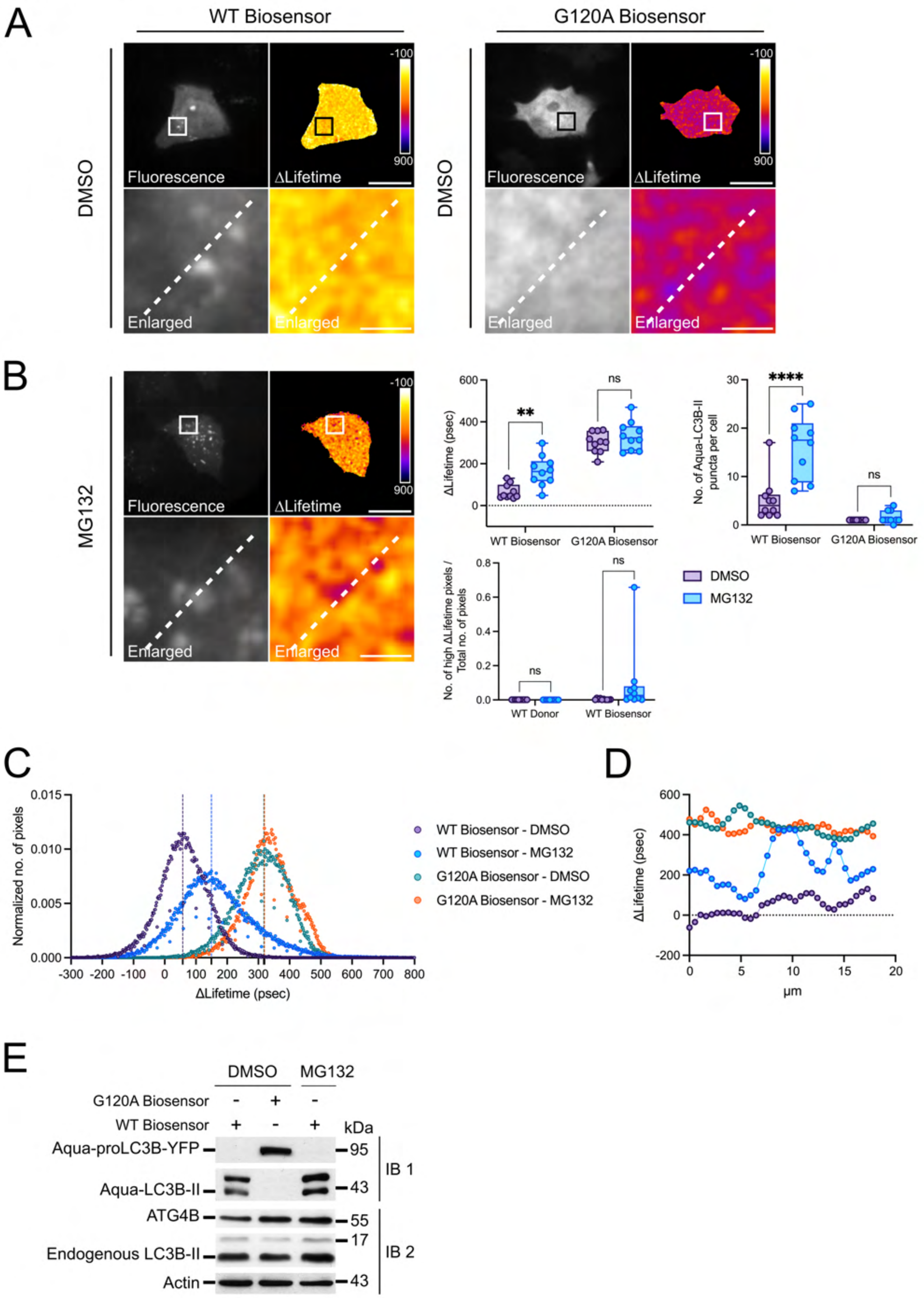
A
WT Biosensor
G120A Biosensor
DMSO
Fluorescence
ΔLifetime
Enlarged
-100
900
B
MG132
ΔLifetime (psec)
WT Biosensor
G120A Biosensor
No. of Aqua-LC3B-II puncta per cell
No. of high ΔLifetime pixels / Total no. of pixels
WT Donor
DMSO
ns
**
****
C
Normalized no. of pixels
WT Biosensor - DMSO
WT Biosensor - MG132
G120A Biosensor - DMSO
G120A Biosensor - MG132
D
µm
E
G120A Biosensor
WT Biosensor
kDa
Aqua-proLC3B-YFP
Aqua-LC3B-II
ATG4B
Endogenous LC3B-II
Actin
IB 1
IB 2
95
43
55
17

### Figure S10

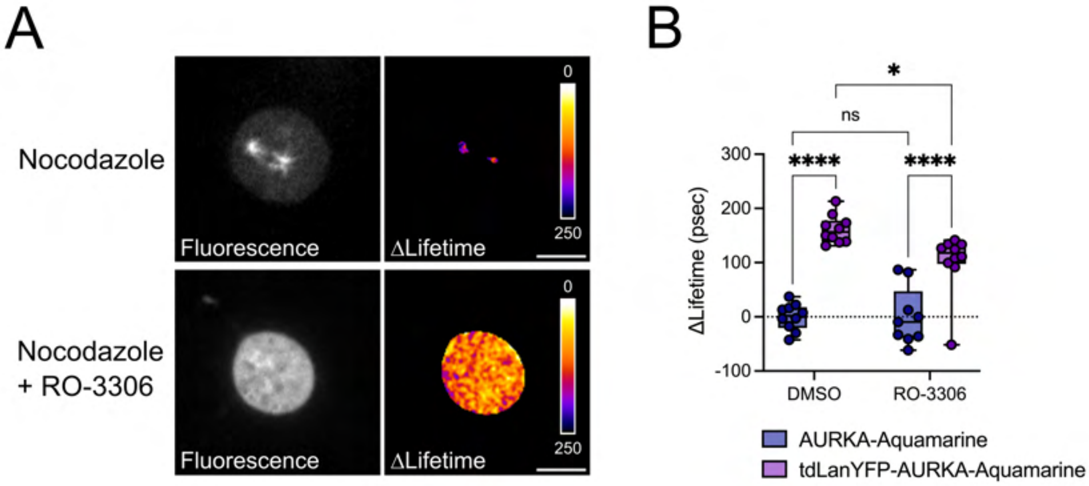
A
Nocodazole
Nocodazole
+ RO-3306
Fluorescence
ΔLifetime
0
250
B
ΔLifetime (psec)
300
200
100
0
-100
DMSO
RO-3306
ns
*
****
AURKA-Aquamarine
tdLanYFP-AURKA-Aquamarine
